# Species-specific regulation of porcine STING stability and antiviral signaling via its K61 mediated K48 ubiquitination and proteasome degradation

**DOI:** 10.64898/2026.03.26.714395

**Authors:** Nengwen Xia, Yajing Chang, Chenglin Chi, Ziyan Sun, Anjing Liu, Wangli Zheng, Jun Jiao, Hongjian Han, Jingyu He, Jiajia Zhang, Nanhua Chen, Sen Jiang, Wanglong Zheng, Jianzhong Zhu

## Abstract

The cGAS-STING pathway has been widely recognized as a critical DNA-sensing pathway that plays a broad-spectrum antiviral role. Livestock, especially pigs, represents one of the most important meat sources. In this study, we identified a key lysine 61 (K61) of porcine STING (pSTING) that plays an essential role in its degradation and antiviral signaling in a species-specific manner, with K61 as the major lysine of pSTING for K48-linked ubiquitination. After virus infection, pSTING recruits the E3 ligase, RNF5, which specifically assembles a K48-linked ubiquitin chain at K61, thereby mediating pSTING proteasomal degradation and reducing its antiviral activity. Meanwhile, the deubiquitylation of K61 is mediated mainly by deubiquitinase USP20, which enhances the stability and antiviral activity of pSTING. Together, given the relatively few lysine numbers in livestock STINGs and species-specific K61 regulation of pSTING stability and antiviral function, the K61 and its specific regulatory enzymes of pSTING could serve as potential targets for breeding of antiviral pigs and design of antiviral drugs, respectively.

## 1. Introduction

Innate immunity serves as the body’s first line of defense against pathogenic microorganisms. It recognizes pathogen-associated molecular patterns (PAMPs), which are common and conserved components of pathogens, as well as damage-associated molecular patterns (DAMPs) derived from the host. This recognition triggers innate immune responses and induces the production of various cytokines, including interferons (IFNs) and inflammatory cytokines, thereby exerting anti-infection effects[1]. The cGAS-STING pathway, a crucial cytosolic DNA-sensing signaling pathway, is involved in innate immune responses elicited by viral and bacterial infections, as well as tissue damage. It serves as a fundamental mechanism for initiating subsequent adaptive immunity. In addition to activating the type I IFN pathway to combat pathogen infection, the cGAS-STING signaling pathway also participates in regulating various biological processes such as autophagy, cell death, DNA damage response, and anti-tumor immunity[2].

Post-translational modifications (PTMs) of proteins include phosphorylation, glycosylation, methylation, acetylation, and ubiquitination, among others. These modifications can alter protein structure, stability, activity, subcellular localization, and interactions with other proteins or small molecular compounds[3, 4]. Ubiquitination is one of the earliest discovered and most extensively studied types of PTMs[5]. Catalyzed by a series of enzymes, ubiquitins are covalently attached to specific lysine (K) residues on target proteins. Depending on the type of ubiquitination or the topology of the ubiquitin chain, the modified target protein may be directed toward distinct functional outcomes or fates. Ubiquitination is involved in diverse biological processes, including the regulation of protein activity and function, protein degradation, cell cycle progression, and cellular differentiation[5, 6].

The activation and inactivation of the cGAS-STING pathway involve various post-translational modifications. For the STING protein, palmitoylation at cysteine 91 (Cys91) is particularly critical for its functional regulation[7, 8]. Palmitoylation modifies the hydrophobicity and membrane-binding properties of STING. Unpalmitoylated STING is primarily localized to the endoplasmic reticulum (ER), whereas after palmitoylation, it translocates from the ER to the Golgi apparatus and can trigger non-cGAS-dependent activation[9]. At the Golgi, TBK1 phosphorylates STING at serine 366 (S366), facilitating the recruitment of IRF3 and subsequent activation of type I IFN signaling[10]. Following activation, STING undergoes degradation via autophagy[11, 12], recycling endosomes[13, 14] or the ubiquitin-proteasome pathway[15, 16] to prevent sustained immune activation. Ubiquitination plays diverse roles in the regulation of STING. In both humans and mice, multiple E3 ligases mediate K48-linked ubiquitination of STING, leading to its proteasomal degradation[15–17]. Additionally, K63-linked polyubiquitination of STING mediates its binding to autophagy receptors and targets it to autolysosomes and recycling endosomes[13, 18, 19], also contributing to negative feedback regulation. On the other hand, K63-linked ubiquitination is essential for the type I IFN activity mediated by STING. Several E3 ligases, including RNF115, TRIM56, and TRIM10 promote STING translocation to the Golgi, dimerization, and subsequent antiviral signaling by enhancing K63-linked ubiquitination[20–22]. In a word, ubiquitination modifications play complex and multifaceted regulatory roles in the various signaling processes mediated by STING.

Here, we identified a species-specific lysine residue at K61 in porcine STING (pSTING) that regulates its protein stability via the K48-linked polyubiquitination. Although the K61 is relatively conserved among mammals, the same polyubiquitination was not observed at the homologous sites in STING proteins from monkey, mouse, cattle or other species. We further identified that K61 is the major site of pSTING for K48-linked ubiquitination and RNF5 is the specifically responsible E3 ubiquitin ligase, which significantly suppresses pSTING protein expression and impairs downstream antiviral signaling. In addition, USP20 acts as a major deubiquitinase for pSTING by removing polyubiquitin chains from K61, thereby stabilizing pSTING protein levels and enhancing pSTING-mediated antiviral responses. In summary, our findings reveal a porcine-specific ubiquitination modification that regulates innate immune signaling, providing not only new insights into the molecular basis of species-specific differences in immune regulation, but also potential targets for breeding of antiviral pigs and design of antiviral drugs.

## 2. Results

### 2.1. Upon activation, porcine STING is degraded through multiple pathways

We first determined that in both porcine primary alveolar macrophages (PAMs) and the porcine alveolar macrophage cell line 3D4/21, stimulation with the double-stranded DNA mimic poly dA:dT but not the RNA mimic poly I:C significantly induced the degradation of STING upon activation and subsequently inhibited the phosphorylation of TBK1 and IRF3 (Figs. 1A and S2A). Notably, only DNA mimic stimulation, not RNA mimic, markedly induced LC3 lipidation and promoted autophagy, which reversely correlated with STING protein expression (Fig. 1A). Further analysis of the degradation kinetics of porcine STING (pSTING) after activation revealed that, whether stimulated with the STING agonist 2’3’-cGAMP in 3D4/21 cells (Fig. 1B) or overexpressed in 293T cells (Fig. 1C), pSTING underwent significant degradation as early as 2 h after activation or under treatment with the translation inhibitor cycloheximide (CHX), eventually decreasing to one-third of its initial levels. The phosphorylation levels of TBK1 also corresponded with the pSTING expression curves (Figs. 1B and 1C).

**Figure 1.**
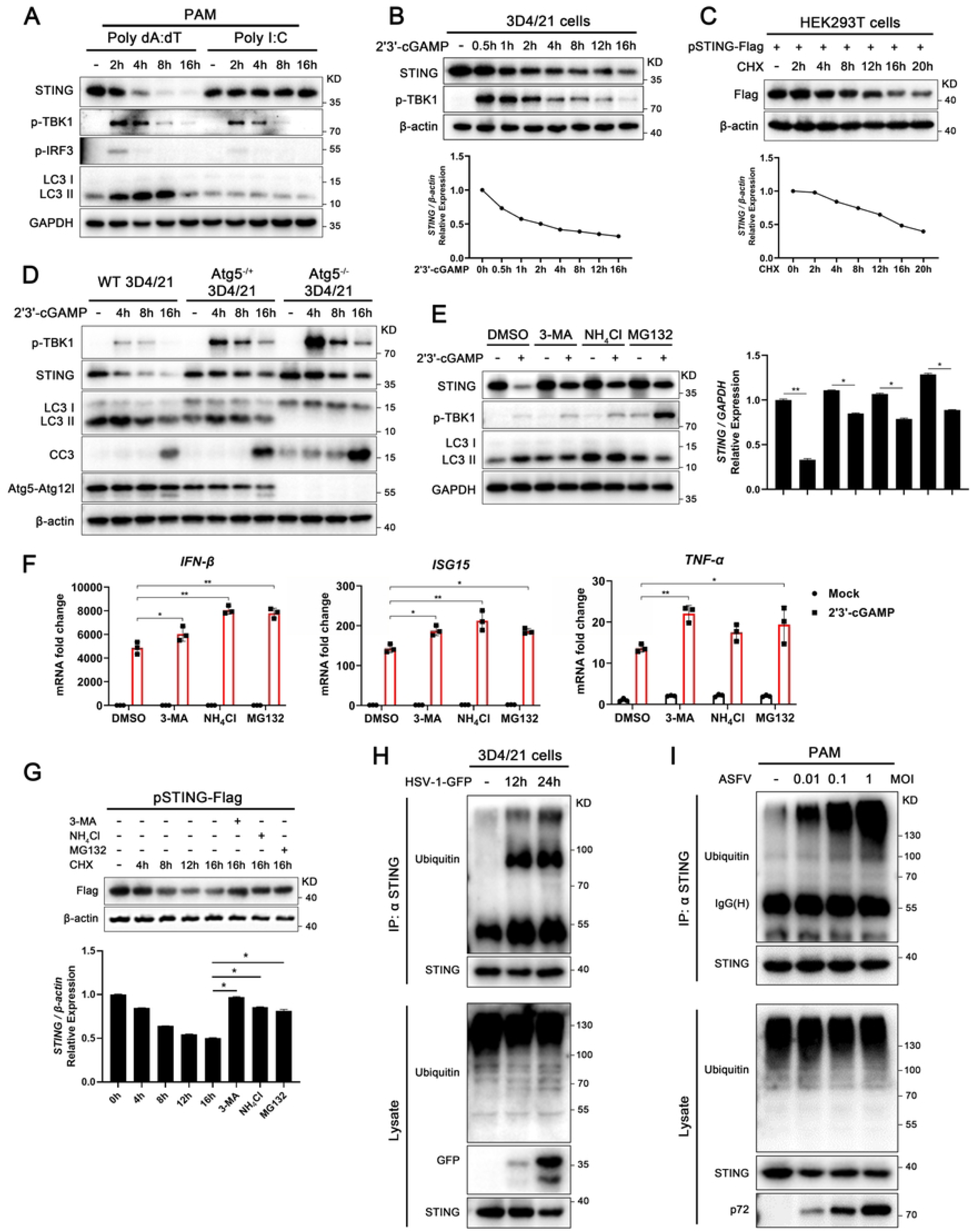
Upon activation, porcine STING is degraded through multiple pathways. **(A)** Primary porcine alveolar macrophages (PAMs) were stimulated by poly dA : dT (2 μg/mL) or poly I:C (2 μg/mL) for the indicated times, cells were harvested and analyzed by Western blotting. **(B-C)** 3D4/21 cells were stimulated by 2’3’-cGAMP (2 μg/mL) for the indicated times (B), HEK293T cells were transfected with Flag-pSTING (1 μg) for 24 h, treated with CHX for the indicated times (C). Cells were then harvested and analyzed by Western blotting. **(D)** ATG5^-/-^3D4/21, ATG5^-/+^ 3D4/21 and wild type cells were stimulated by 2’3’-cGAMP (2 μg/mL) for the indicated times, the expression of the target proteins was determined by Western blotting. **(E-G)** 3D4/21 cells were stimulated by 2’3’-cGAMP (2 μg/mL) for 8 h (E and F), HEK293T cells were transfected with Flag-pSTING (1 μg) for 24 h, treated with CHX for the indicated times (G). Prior to harvesting, cells were treated with 3-MA (50 μM), NH_4_Cl (25 mM), and MG132 (10 μM) for 0.5 h, and then then analyzed using RT-qPCR (F) and Western blotting (E and G). **(H-I)** 3D4/21 cells were infected with HSV-1 (MOI=0.1) for indicated times (H), PAMs were infected with ASFV at the specified MOIs for 48h (I), and then cell were treated by MG132 (10 μM) for 6 h. After cell collection, STING was immunoprecipitated, and its ubiquitination level was determined by Western blotting. In the panels B, C, E and G, the expressions of STING or Flag-pSTING were normalized to β-actin or GAPDH, and the densitometry values were plotted. \**p* < 0.05 and \*\**p* < 0.01.

Changes in the conformation and cytoplasmic localization of STING during activation trigger both canonical and non-canonical autophagy via an interferon-independent pathway[11, 23, 24], which serves as a necessary negative feedback mechanism to prevent excessive STING activation. Our results also confirmed that pSTING degradation is highly consistent with the induction of autophagy (Fig. 1A). However, in our further investigations, we found that impaired autophagy resulting from ATG5 knockdown/deficiency did not completely inhibit pSTING degradation induced by its activation (Fig. 1D and Fig. S1A and B). Similarly, in both 3D4/21 and 293T cells, treatment with 3-MA, NH_4_Cl, or MG132, which inhibit autophagy, lysosomal function, and proteasome activity, respectively, all partially reversed pSTING degradation after activation and enhanced multiple downstream signaling cascades, including autophagy and type I interferon (IFN) responses (Figs. 1E–1G). Therefore, we concluded that, in addition to the autophagy-lysosome pathway, the ubiquitin-proteasome pathway is also essential for the negative regulation of pSTING stability and signaling. Consistently, in 3D4/21 cells, infection with HSV-1 or stimulation with 2’3’-cGAMP revealed that, over time, activated pSTING underwent pronounced polyubiquitination (Figs. 1H and S2B). Furthermore, similar results were observed in porcine primary alveolar macrophages (PAM) infected with African swine fever virus (ASFV), with the ubiquitination level of STING heightened upon increase of ASFV infection (Fig. 1I). These results demonstrated that after DNA virus infection, activated pSTING is degraded primarily through both the autophagy-lysosome and ubiquitin-proteasome pathways.

### 2.2. The lysine 61 is crucial for the stability of porcine STING protein and its antiviral activity

STING-mediated autophagy is an ancient and conserved function that has been extensively studied in recent years[11, 25]. While the ubiquitin-proteasome degradation pathway of STING has been well characterized in humans and mice, its mechanism remains poorly defined in pigs. Moreover, reported findings often vary across species, prompting us to investigate the mechanism mediating ubiquitin-proteasome degradation of porcine STING (pSTING). Since ubiquitination typically occurs on lysine (K) residues of target proteins, we individually mutated each of the six lysines into arginines (K–R) on pSTING, generated six mutants each retaining only one lysine while mutating the other five into arginines (K–O), and created a mutant with all six lysines replaced by arginines (K0), resulting in 13 STING mutants in total. Luciferase reporter assays revealed that mutation of the K61R significantly enhanced pSTING protein expression and increased the activity of downstream ISRE and NF-κB promoters (Fig. 2A). Conversely, the pSTING K61O mutant, which retains only K61 while mutating the other five K, exhibited protein expression and downstream promoter activity comparable to wild-type pSTING (Fig. 2B). This indicated that the K61 site is critical for regulating pSTING protein stability.

**Figure 2.**
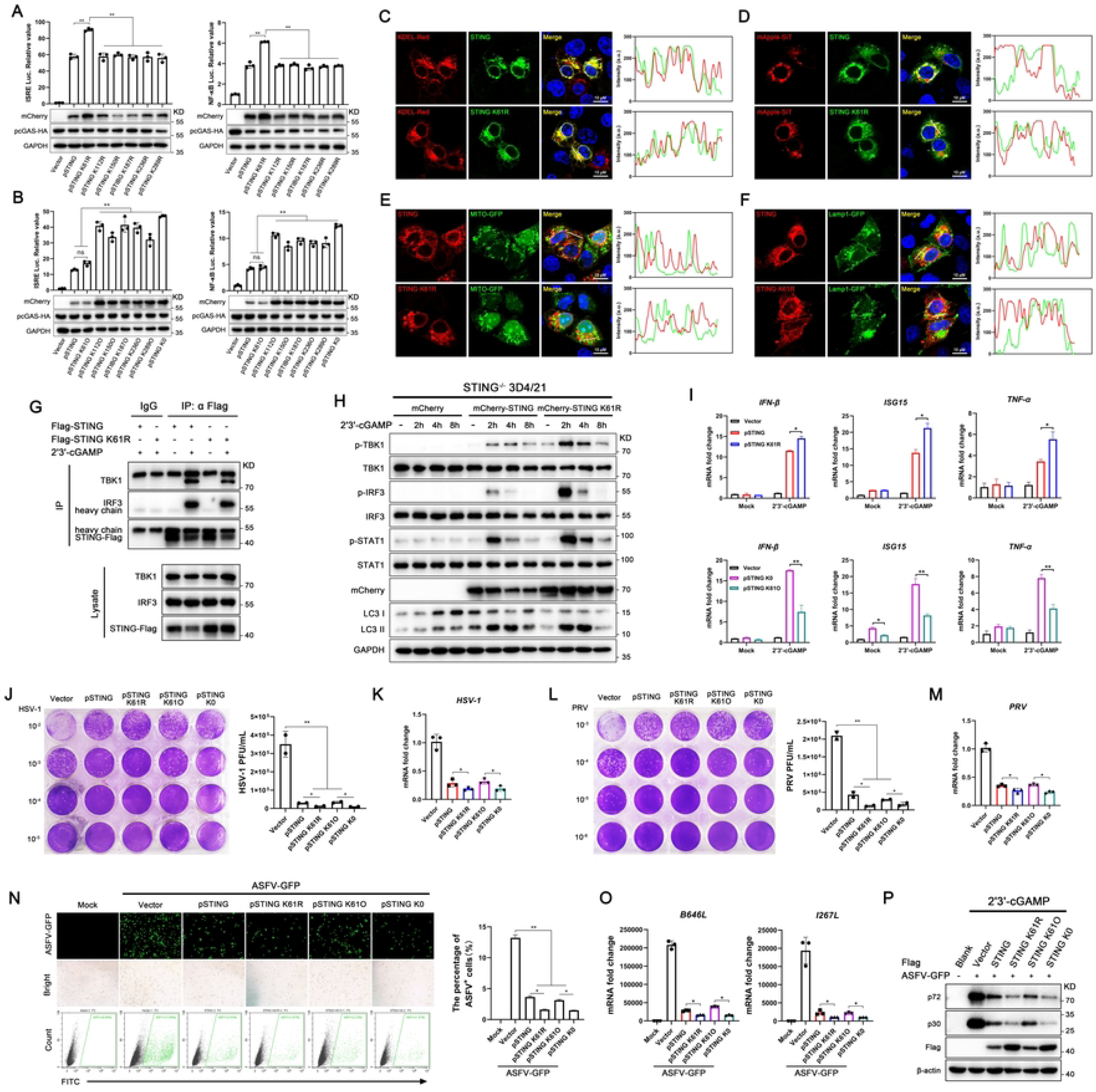
The lysine 61 is crucial for the stability of porcine STING protein. **(A-B)** HEK293T cells were co-transfected with mCherry-pSTING (20 ng) or its mutants together with pcGAS-HA (20 ng), plus ISRE-luc (10 ng) or NF-κΒ-luc (10 ng) and pRL-TK plasmid (0.4 ng) for 24 h. Luciferase activities were detected, and protein expressions were analyzed. **(C-F)** 3D4/21 cells in 24-well cell crawl slides were co-transfected with Flag-pSTING or K61R mutant together with KDEL-Red (C), mApple-SiT (D), MITO-GFP (E) or Lamp1-GFP (F) (each 0.5 μg) for 24 h, and then cells were fixed, stained and the colocalization of STING and its K61R mutant with organelles was examined by confocal fluorescence microscopy. **(G)** HEK293T cells in 6-well were transfected with Flag-pSTING or K61R mutant (1 μg/mL) for 24h, and stimulated with 2’3’-cGAMP (2 μg/mL) for 2 h. Then interaction between STING and TBK1/IRF3 was identified by Co-IP using anti-FLAG antibody, whereas rabbit IgG served as the control. **(H-I)** STING^-/-^ 3D4/21 cells in 12-well were transfected with pmCherry-C1, mCherry-pSTING or K61R mutant (1 μg) for 24 h, and stimulated with 2’3’-cGAMP (2 μg/mL) for indicated times (H) or 6 h (I). Cells were harvested and analyzed by Western blotting (H) and RT-qPCR (I). **(J-M)** STING^-/-^ 3D4/21 cells were transfected with Flag-pSTING or its mutants (1μg), and then stimulated with 2’3’-cGAMP (2 μg/mL) for 2 h, followed by infections with HSV-1 (MOI=0.01, 36 h) or PRV (MOI=0.01, 24 h). The viral titers in the supernatants from HSV-1 (J) or PRV (L) infected cells was measured by plaque assay, and the HSV-1 and PRV gD gene expressions were measured by RT-qPCR (K and M). (**N-P**) STING^-/-^ MA104 cells were transfected and stimulated as in STING^-/-^ 3D4/21 cells, followed by infections ASFV-GFP, infected cells were detected by fluorescence microscopy and cytometry, Western blotting and quantitative PCR. \**p* < 0.05 and \*\**p* < 0.01.

To further determine whether the K61 affects pSTING degradation itself rather than its transport from the endoplasmic reticulum (ER) to the Golgi apparatus, we co-expressed pSTING and pSTING K61R with organelle-specific markers for the ER, Golgi, mitochondria, and lysosomes in 3D4/21 cells. The results showed that mutation of K61R did not alter the localization of pSTING to the ER and Golgi (Figs. 2C and 2D). And pSTING showed no association with mitochondria regardless of the presence of K61 (Fig. 2E), with the K61R mutation increasing pSTING accumulation in lysosomes to certain degree (Fig. 2F). Furthermore, the K61R mutation in pSTING did not affect the recruitment of TBK1 and IRF3 following stimulation with a STING agonist (Fig. 2G), indicating that the presence of the K61 is not essential for the pSTING activation process.

We next transiently expressed pSTING and related mutants in STING^-/-^ 3D4/21 cells and HEK293T cells using transfection to examine alterations in downstream signaling. Consistent with the promoter activity results, Western blot analyses and qPCR demonstrated that the mutation of K61R significantly enhanced STING activation-induced phosphorylations of TBK1 and IRF3 and the downstream gene transcriptions (Figs. 2H–2I and S2C–S2D). In addition, the K61R mutation markedly enhanced LC3 lipidation after stimulation with the STING agonist 2’3’-cGAMP (Fig. 2H), indicating abnormally elevated autophagy levels, which also explains the higher lysosome accumulation observed with the K61R mutant (Fig. 2F). Since cGAS-STING signaling activation can induce multiple types of cell death[26], including apoptosis[27], we used flow cytometry to assess the effect of STING and its mutants on apoptosis. The results showed that apoptosis induced by STING activation remained low when K61 was present, while mutation of K61R alone maximally enhanced pSTING-mediated apoptosis (Figs. S2E and S2F). Treatment with MG132 revealed that proteasome inhibition obviously alleviated the increase in STING expression caused by the K61R mutation, indicating that the K61 site is closely associated with proteasome-mediated degradation of porcine STING (Fig. S2G).

We further evaluated the antiviral capacity of STING with K61 retained or mutated by complementing STING^-/-^ 3D4/21 cells or STING^-/-^ MA104 cells with STING and related mutants, followed by infection with three DNA viruses (HSV-1, PRV and ASFV) and an RNA virus VSV. The viral titration results demonstrated that mutation of the K61 site significantly enhanced STING-mediated antiviral activities against HSV-1, PRV, ASFV and VSV infections in plaque assay (HSV-1, PRV and VSV), viral gene qPCR quantification assay (HSV-1, PRV and ASFV) and viral protein expression quantification assay (ASFV) (Figs. 2J–2P and S2H-S2I).

### 2.3. The K61 site of porcine STING is the predominant site for K48-linked polyubiquitination

Since K61 acts as a negative regulatory site in the modulation of STING protein stability and is closely associated with proteasomal activity, we wondered whether and what kind of polyubiquitination modification at this site mediates the degradation of the STING. To investigate whether ubiquitin molecules are attached to this site and to determine the specific linkage types, we co-transfected 293T cells with pSTING or pSTING K61R along with eight types of ubiquitin expression plasmids, including wild-type ubiquitin (Fig 3A). Our results demonstrated that pSTING can undergo ubiquitination with multiple linkage types; however, the K61R mutation significantly suppresses K27-, K29-, K33- or K48-linked polyubiquitination (Figs. 3A and 3B), suggesting that K61 site of pSTING is subjected for K27-, K29-, K33- and K48-linked polyubiquitination.

**Figure 3.**
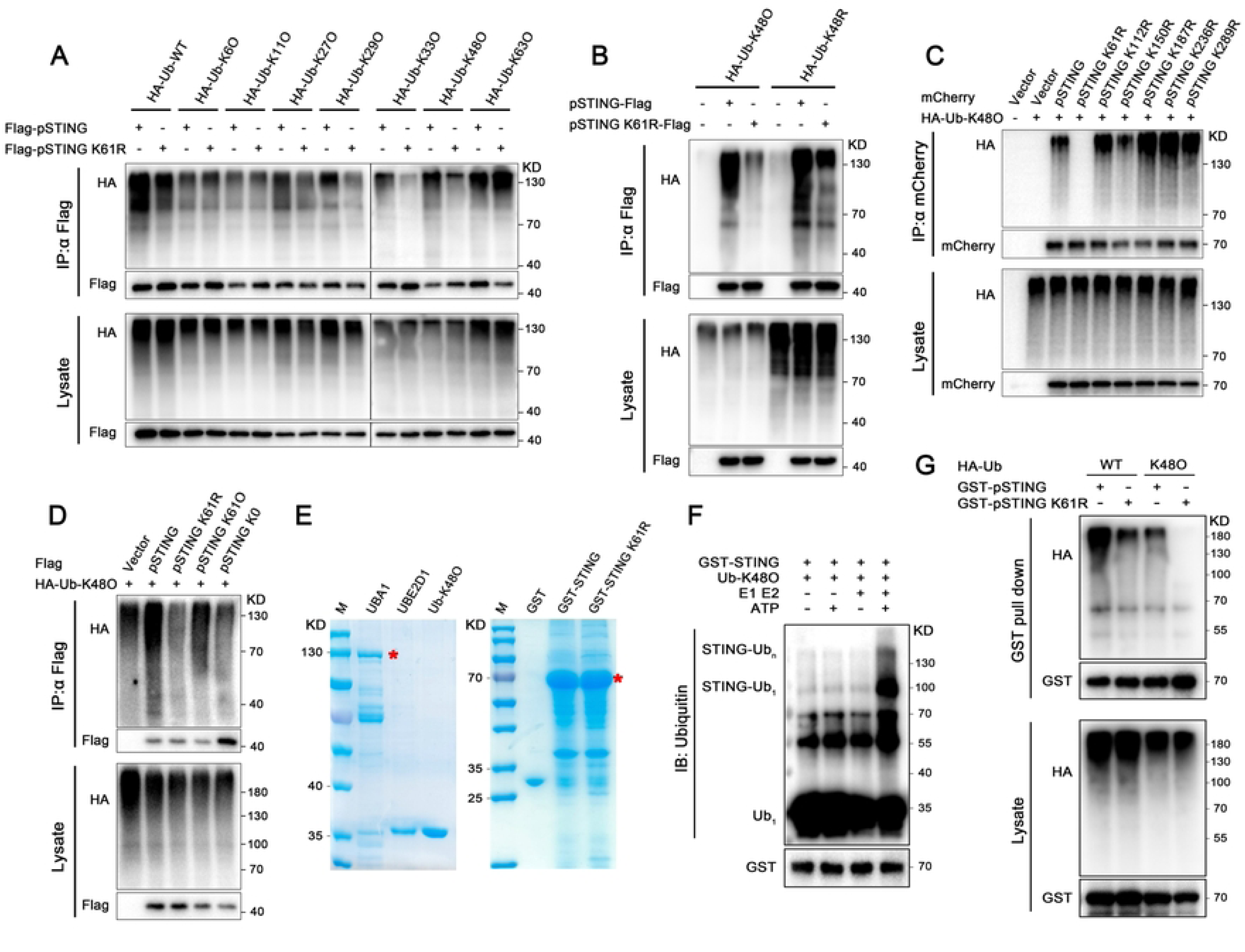
Porcine STING is ubiquitinated on lysine (K) 61 with K48-linked polyubiquitin chains. **(A-D)** HEK293T cells in 6-well were co-transfected with Flag-pSTING or its mutants and HA-Ub-WT or its mutants (each 0.5 μg) for 24 h, and then transfected cells were treated with MG132 (10 μM) for 6 h. STING was immunoprecipitated, and its ubiquitination levels were determined by Western blotting. (**E**) Bacterial expressed and purified UBA1, UBE2D1, Ub-K48O, GST, GST-STING and GST-STING K61R proteins were analyzed by SDS-PAGE and Coomassie blue staining. (**F**) The specified purified proteins were incubated at 37°C for 2 h in the presence or absence of ATP, heat-inactivated at 95°C for 10 min, and the ubiquitination level of STING was analyzed by Western blotting. (**G**) The purified GST-pSTING or the K61R mutant was incubated with lysates of HEK293T cells transfected with HA-Ub-WT or HA-Ub-K48O at 37°C for 2 h. Following GST pull-down, the ubiquitination level of STING was detected by Western blotting.

Since K48-linked polyubiquitination is a key signal for proteasomal degradation, we further examined the impact of other K residues on K48-linked polyubiquitination of pSTING. We found that mutation of K61R alone almost completely abolishes K48-linked polyubiquitination of pSTING, despite that mutation of K150R also slightly reduces it (Fig. 3C). By incorporating STING K61O and K0 mutants, we further characterized the polyubiquitination patterns at K61 site. The results showed that the K61R mutation nearly eliminates all K48-linked ubiquitination of pSTING (Fig. 3D) but only attenuates K27-, K29- and K33-linked ubiquitination (Figs. S3A–S3C). On the other hand, keeping K61 alone of K61O mutation almost exhibited same level of K48-linked ubiquitination as wide type pSTING, but insufficient to restore full K27-, K29-, or K33-linked ubiquitination in pSTING (Fig 3D and Figs. S3A–S3C). These results clearly suggested that the K61 is the major site of K48-linked ubiquitination of pSTING, whereas multiple K sites of pSTING including K61 have K27-, K29- and K33-linked ubiquitinations.

By expressing and purifying porcine UBA1 (E1 activating enzyme), UBE2D1 (E2 conjugating enzyme), Ub-K48, and GST-tagged pSTING and K61R mutant in *E. coli* (Fig. 3E), we found that pSTING undergoes K48-linked ubiquitination in the presence of E1, E2 and ATP with *in vitro* ubiquitination assay (Fig. 3F). Further, cell lysates of 293T cells transfected with Ub-WT and Ub-K48O were incubated with purified GST-pSTING and GST-pSTING K61R, respectively, and the results demonstrated that the K61 site is essential for K48-linked ubiquitination modification of pSTING (Fig. 3G).

### 2.4. The ubiquitination of porcine STING at K61 regulates its protein stability in a species-specific manner

It is unknown that whether the K61 site is conversed and crucial for the stability of STING protein in other species. To investigate this, we conducted a comparative analysis of the conservation of this site across STING proteins from nine species, including human, monkey, mouse, pig, cattle and avian species. The results showed that the K61 homologous site is highly conserved in other mammals and even in avian species with the exception of human (Fig. 4A). This led us to hypothesize that the homologous K61 site in other species might possess a similar function. We subsequently cloned the STING genes from monkey (mo), mouse (m), bovine (b), and chicken (ch), along with K to R mutants of their respective homologous K61 sites. These various STING genes were expressed in transfected 293T cells to examine steady-state STING protein levels and assay downstream signaling (Figs. 4B-D). Contrary to our expectations, mutations of K to R at the homologous K61 sites in species other than pigs did not affect their protein stability, nor did they enhance the subsequent phosphorylation of TBK1 and IRF3 in Western blotting (WB) (Fig. 4B). Results from the dual-luciferase reporter assay and qRT-PCR were consistent with the WB results, that is, only the mutation of pSTING K61R significantly potentiated the downstream type I IFN signaling (Figs. 4C and 4D). We further assessed the ubiquitin modification of the STING K61 homologous site mutants from these species (Figs. 4E-I). The results indicated that only the pSTING K61R mutation significantly reduced the levels of total ubiquitination and K48-linked ubiquitination (Fig. 4E). In contrast, the K to R mutations at the homologous K61 sites in moSTING, mSTING, bSTING and chSTING had no effect on the ubiquitin modifications of these STING proteins (Figs. 4F–4I). These results clearly demonstrated that the K48-linked polyubiquitination on the K61 site of pSTING is species-specific and unique.

**Figure 4.**
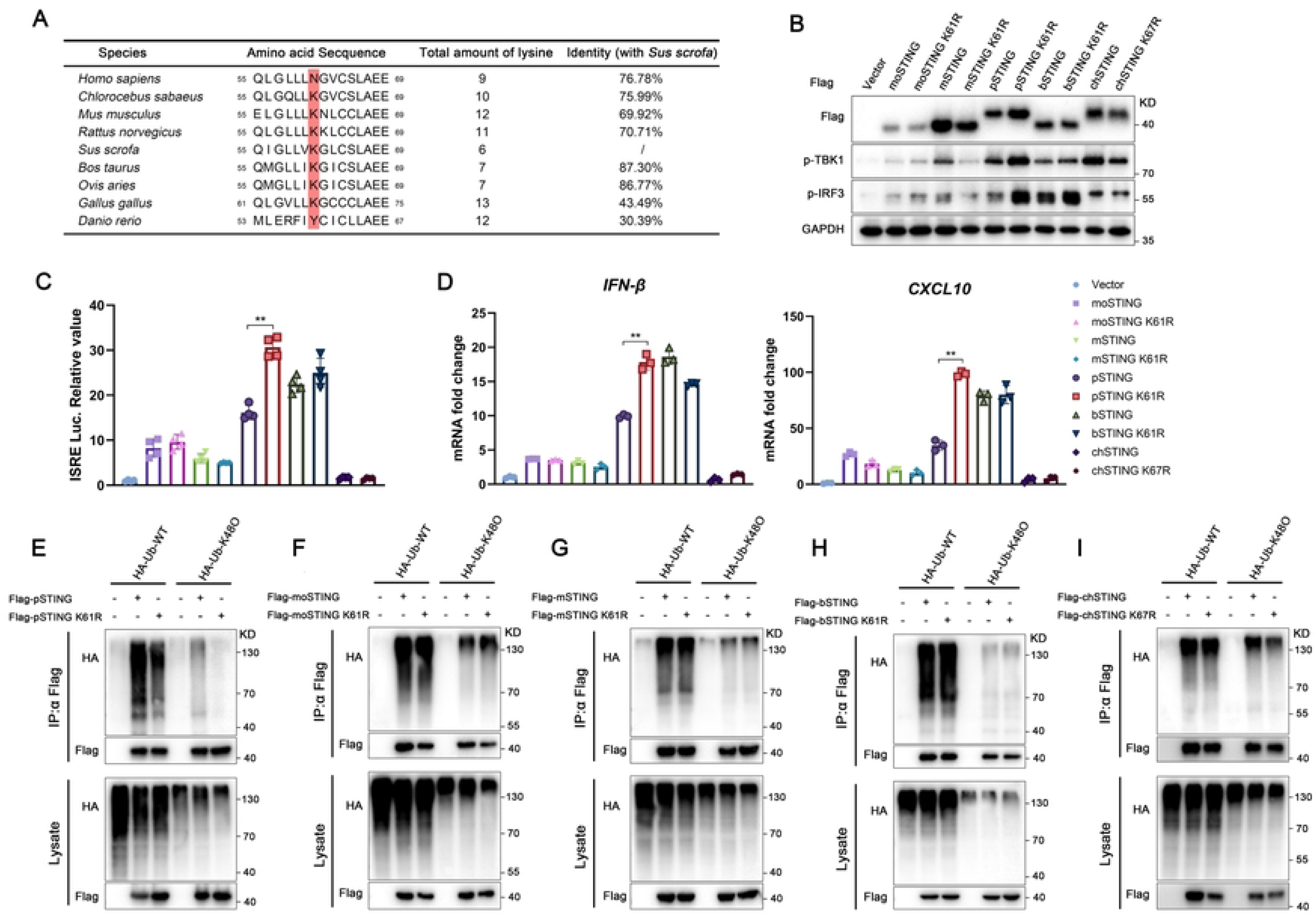
The ubiquitination of porcine STING at K61 regulates its protein stability in a species-specific manner. **(A)** Sequence alignment of STING K61 or the homologous amino acids (highlighted in red) from human, monkey, mouse, rat, pig, cattle, sheep, chicken and zebrafish. **(B-D)** HEK293T cells in 12-well were co-transfected with Flag-tagged STING and K61 homologous site mutants from various species (each 1 μg) plus ISRE-luc (200 ng) and pRL-TK plasmid (10 ng) for 24 h. Cells were harvested and analyzed by Western blotting (B), luciferase assay (C) and RT-qPCR (D). **(E-I)** HEK293T cells in 6-well were co-transfected with Flag-tagged porcine (p) STING (E), monkey (mo) STING (F), mouse (m) STING (G), bovine (b) STING (H), chicken (ch) STING (I) and K61 homologous site mutants from various species, plus HA-Ub-WT or HA-Ub-K48O (each 0.5 μg) for 24 h, and then cells were treated by MG132 (10 μM) for 6 h. STING was immunoprecipitated, and its ubiquitination levels were determined by Western blotting. \**p* < 0.05 and \*\**p* < 0.01.

Unlike other types of mammals, various livestock species, including pig, cattle, and sheep, possess STING proteins with fewer lysine (K) residues, suggesting potential species-specific differences (Fig. 4A). The homology between bovine STING and porcine STING is 87.3%, which is significantly higher than the homology between porcine STING and STING from other non-livestock species (Fig. 4A). This led us to question why the K61 site in bovine STING does not share a similar function. We hypothesized that differences in the amino acids flanking the K61 site in the STING proteins of these two species might affect ubiquitin binding. By comparing the amino acid sequences around the K61 site in both species, we generated a bovine STING mutant (named M1) containing the I60V and I63L mutations, making these positions identical to those in porcine STING (V60 and L63) (Fig. S3D). We also created a bovine STING mutant (named M2) with the I60V, K61R, and I63L mutations (Fig. S3D). Subsequently, we examined the ubiquitination profiles of these different bovine STING mutants. Surprisingly, and consistent with our hypothesis, mutating the amino acids near the K61 site (M1) enhanced both the total ubiquitination and K48-linked polyubiquitination of bovine STING. Further, when the K61R mutation was also introduced (M2), this enhancing effect was significantly inhibited (Figs. S3E and S3F).

### 2.5. RNF5 acts as an E3 ligase to promote K29-, K33- and K48-linked polyubiquitination of porcine STING and specifically K48-linked polyubiquitination of K61

The process by which ubiquitin molecules bind to target proteins and modify them requires the participation of three types of ubiquitin-related enzymes. Among these, E3 ubiquitin ligases transfer the activated ubiquitin molecule from the E2 conjugating enzyme to the substrate protein, thereby achieving ubiquitination of the target protein[28]. Unlike the relatively small number of E1 activating enzymes and E2 conjugating enzymes[29], there are hundreds of E3 ubiquitin ligases in mammals[30]. To identify the E3 ubiquitin ligase mediating polyubiquitination at the K61 site of STING in pigs, we cloned several E3 ligases reported to mediate STING degradation in humans and mice, which include TRIM29, TRIM13, RNF90, and RNF5. Among them, TRIM29, RNF90, and RNF5 promote the ubiquitin-proteasome degradation pathway of STING by enhancing its K48-linked polyubiquitination[15–17], while TRIM13 catalyzes Lys63-linked polyubiquitination of STING, leading to slowed ER exit and accelerated ER-associated degradation of STING[31].

By co-expressions of the porcine E3 ligases with pSTING protein in 293T cells, we found that only co-expression with RNF5 significantly reduced pSTING protein levels (Fig. 5A). Further mechanistic analysis revealed that, despite that the four E3 ligases all show clear interaction and cytoplasmic co-localization with pSTING, only RNF5 and RNF90 significantly enhance both total ubiquitination and K48-linked polyubiquitination of STING (Figs. 5B–5D and S4A-S4C). Co-immunoprecipitation of endogenous STING in 3D3/21 cells demonstrated that HSV-1 infection markedly strengthens the interaction between STING and RNF5 (Fig. 5E), a trend consistent with the ubiquitination level of STING post virus infection (Figs. 1H). Furthermore, RNF5 inhibited various downstream signals of STING, including IFN response and autophagy, in a dose-dependent manner by promoting STING degradation (Figs. 5F and S4D and S4E). Notably, the K61R mutation of pSTING significantly reversed the inhibitory effect of RNF5 on pSTING degradation and downstream signaling transduction (Figs. S4F and S4G). Through expression and purification of the porcine RNF5 protein in *E. coli* (Fig. 5G), GST pull-down and in *vitro* ubiquitination assays demonstrated that porcine RNF5 can interact with pSTING directly (Fig. 5H) and mediate K48-linked ubiquitination of pSTING (Fig. 5I). We also did the expression and purification of K48-linked polyubiquitin chains (composed of four K48-linked ubiquitin units, Ub_4_) (Fig. 5G). Consistently, RNF5 significantly enhanced K48-linked pSTING monoubiquitination (Fig 5I) as well as polyubiquitination (Fig. 5J) in the *in vitro* ubiquitination assay. Moreover, RNF5 also significantly enhanced K29-, K33-linked polyubiquitination in addition to K48-linked polyubiquitination of pSTING (Fig. 5K). When K61R mutation occurred, RNF5 significantly enhanced K29- and K33-linked polyubiquitination of pSTING without promoting K48-linked polyubiquitination (Fig. 5L). Conversely, when K61O mutation occurred, RNF5 specifically enhanced K48-linked polyubiquitination without promoting other linkage types (Fig. 5M). These results indicated that K61 is the critical site mediating K48-linked polyubiquitination of pSTING, and that RNF5, as the E3 ligase, specifically mediates K48-linked polyubiquitination at this site.

**Figure 5.**
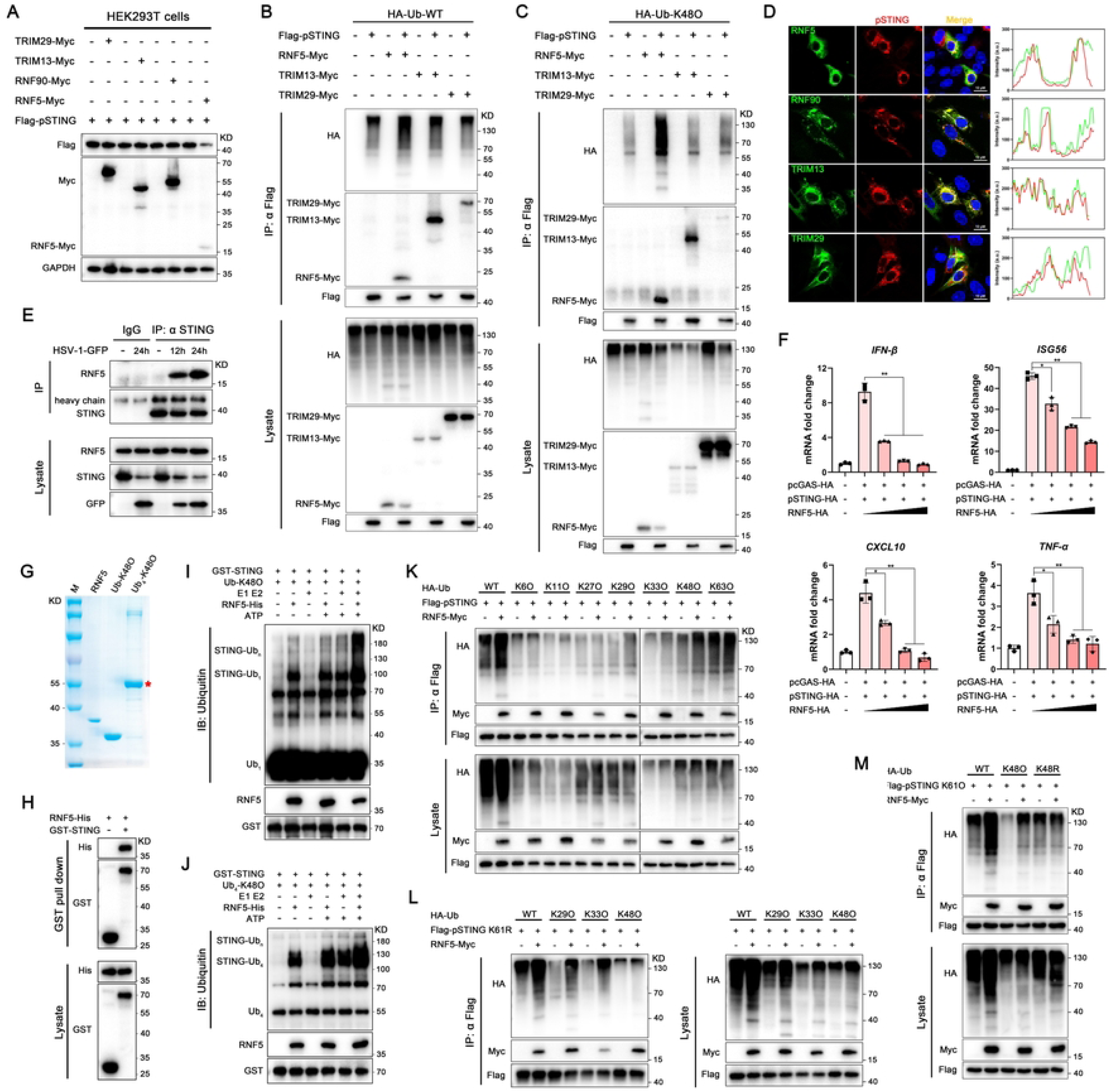
RNF5 acts as an E3 ligase to promote K29-, K33- and K48-linked polyubiquitination of pSTING and K48-polyubiquitinaiton specifically on K61. **(A)** HEK293T cells were co-transfected with Flag-pSTING and TRIM29-Myc, TRIM13-Myc, RNF90-Myc or RNF5-Myc (each 0.5 μg) for 24 h, and then the expressions of pSTING were determined by Western blotting. **(B** and **C)** HEK293T cells were co-transfected with Flag-pSTING and RNF5-Myc, TRIM13-Myc or TRIM29-Myc plus HA-Ub-WT (B) or HA-Ub-K48O (C) (each 0.7 μg) for 24 h, and then cells were treated by MG132 (10 μM) for 6 h. STING was immunoprecipitated, and its ubiquitination levels were determined by Western blotting. **(D)** 3D4/21 cells in 24-well cell crawl slides were co-transfected with Flag-pSTING and RNF5-Myc, TRIM13-Myc, TRIM29-Myc, or RNF90 (each 0.5 μg) for 24 h. The cells were then fixed, stained and examined by confocal microscopy to assess the colocalization of STING with the respective E3 ligases. **(E)** 3D4/21 cells were infected with HSV-1 for indicated times, then interaction between STING and RNF5 was identified by Co-IP using anti-STING antibody, whereas rabbit IgG served as the control. **(F)** RT-qPCR was used to detect the effects of varying doses of RNF5(0, 100, 200, 400 ng) on the transcription of STING -activated downstream genes in transfected HEK293T cells. **(G)** Bacterial purified RNF5, Ub-K48O and Ub_4_-K48O proteins were analyzed by SDS-PAGE and Coomassie blue staining. **(H)** The interaction between purified GST-STING and RNF5-His *in vitro* was analyzed by GST pull-down assay. **(I and J)** The specified purified proteins together with Ub-K48 (I) or Ub_4_-K48 (J) were incubated at 37°C for 2 h in the presence or absence of ATP and RNF5, heat-inactivated at 95°C for 10 min, and the ubiquitination level of STING was analyzed by Western blotting. **(K-M)** HEK293T cells were co-transfected with RNF5 and pSTING (K), pSTING K61R (L) or pSTING K61O (M), plus HA-Ub-WT or its mutants (each 0.7 μg) for 24 h, and then cells were treated by MG132 (10 μM) for 6 h. STING was immunoprecipitated, and its ubiquitination level was determined by Western blotting. \**p* < 0.05 and \*\**p* < 0.01.

### 2.6. The CBD domain of porcine STING and the TM domain of RNF5 are key regions for their mutual interaction

To identify the critical domains responsible for the RNF5-STING interaction and the subsequent promotion of ubiquitination, we analyzed the key domains of STING and RNF5 using the UniProt online database (https://www.uniprot.org/) and generated corresponding truncation mutants. The STING domains include an N-terminal transmembrane (TM) domain, a central cyclic dinucleotide (cGAMP) binding domain (CBD), and a C-terminal tail (CTT) region that is crucial for regulating type I IFN signaling. By selectively retaining or deleting specific domains, we created eight STING-related truncation/deletion mutants (Fig. 6A). Since amino acids 238-241 of STING were reported to be critical for mediating autophagy[11], we also constructed a mutant lacking this region (Fig. 6A). Reciprocal co-immunoprecipitation (Co-IP) assays revealed that the presence of the CBD domain alone was sufficient to mediate a strong interaction with RNF5, and deletion of this domain completely abolished the interaction (Figs. 6B–6C and S5A). This finding differed from previous reports in human systems[16], indicating the existed species specificity. Intracellular co-localization results further confirmed the findings; deletion of the porcine STING CBD domain significantly altered the protein’s subcellular localization and markedly reduced its co-localization with RNF5 (Fig. 6D).

**Figure 6.**
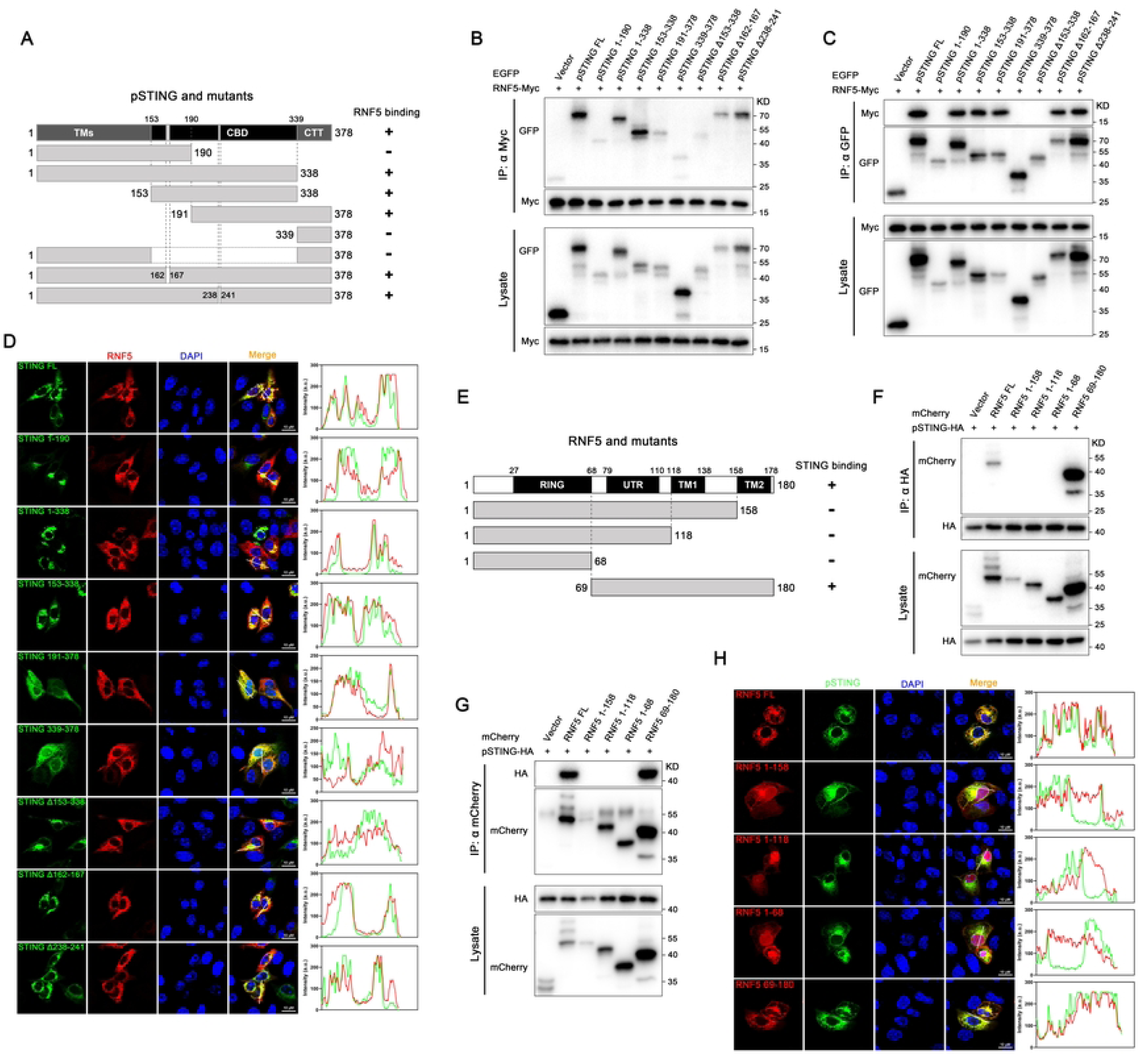
The CBD domain of pSTING and the TM domain of RNF5 are key regions for their mutual interaction. **(A-C)** A schematic diagram of pSTING truncations (A), and reciprocal co-immunoprecipitation analysis of the interaction between pSTING domains and RNF5 in HEK293T cells co-transfected with EGFP-STING truncations and RNF5-Myc (B and C). **(D)** The colocalization of pSTING truncations with pRNF5 was observed by confocal microscopy. **(E-G)** A schematic diagram of RNF5 truncations (E), and reciprocal co-immunoprecipitation analysis of the interaction between RNF5 domains and pSTING in HEK293T cells co-transfected with mCherry-RNF5 truncations and pSTING-HA (F and G). **(H)** The colocalization of pRNF5 truncations with pSTING was observed by confocal microscopy.

RNF5 is a typical E3 ubiquitin ligase containing a RING finger domain and is a small, structurally simple protein. Besides the RING domain, it also contains a domain of undefined function and two transmembrane regions (TM1 and TM2) (Fig. 6E). We systematically deleted segments of porcine RNF5 and assessed the impact on its interaction with pSTING. Reciprocal Co-IP results demonstrated that deletion of the TM2 domain alone significantly inhibited the RNF5-STING interaction, whereas deletion of the RING domain had no effect (Figs. 6F–6G and S5B). Immunofluorescence analysis further showed that full-length RNF5 is exclusively cytoplasmic and shows strong co-localization with pSTING; however, deletion of TM2 alters RNF5’s subcellular localization, causing it to accumulate predominantly in the nucleus and drastically reducing its co-localization with STING (Fig. 6H). Consistent with the protein interaction, deletion of the TM2 domain of RNF5 not only abolished its interaction with pSTING, but also eliminated its ability to promote pSTING ubiquitination (Fig. S5C).

### 2.7. RNF5 negatively regulates porcine STING-mediated antiviral response

To further elucidate the regulatory role of RNF5 on STING and downstream signaling, we generated RNF5-knockout homozygous clones in 3D4/21 cells using CRISPR-Cas9 (Fig. S1D and E). Two homozygous clones were selected to evaluate their responses to STING agonists. Quantitative real-time PCR results demonstrated that RNF5 knockout enhanced the transcription of IFN-β and related interferon-stimulated genes (ISGs) following STING activation (Fig. 7A). Mechanistically, in porcine macrophages, RNF5 knockout significantly increased STING protein expression, delayed its degradation upon activation, and enhanced the phosphorylation of downstream TBK1 and IRF3 (Fig. 7B), thereby potentiating STING-mediated type I IFN responses. When RNF5-deficient cells were treated with the autophagy and lysosome inhibitors 3-MA and NH_4_Cl, simultaneous inhibition of both the ubiquitin-proteasome and autophagy-lysosome pathways maximally attenuated STING degradation upon activation and enhanced downstream signaling (Figs. 7C and 7D). Further investigation into STING ubiquitination in RNF5-knockout cells revealed that, similar to observations in 293T cells, RNF5 knockout significantly reduced STING ubiquitination levels post 2’3’-cGAMP treatment, although not completely abolished (Fig. 7E).

**Figure 7.**
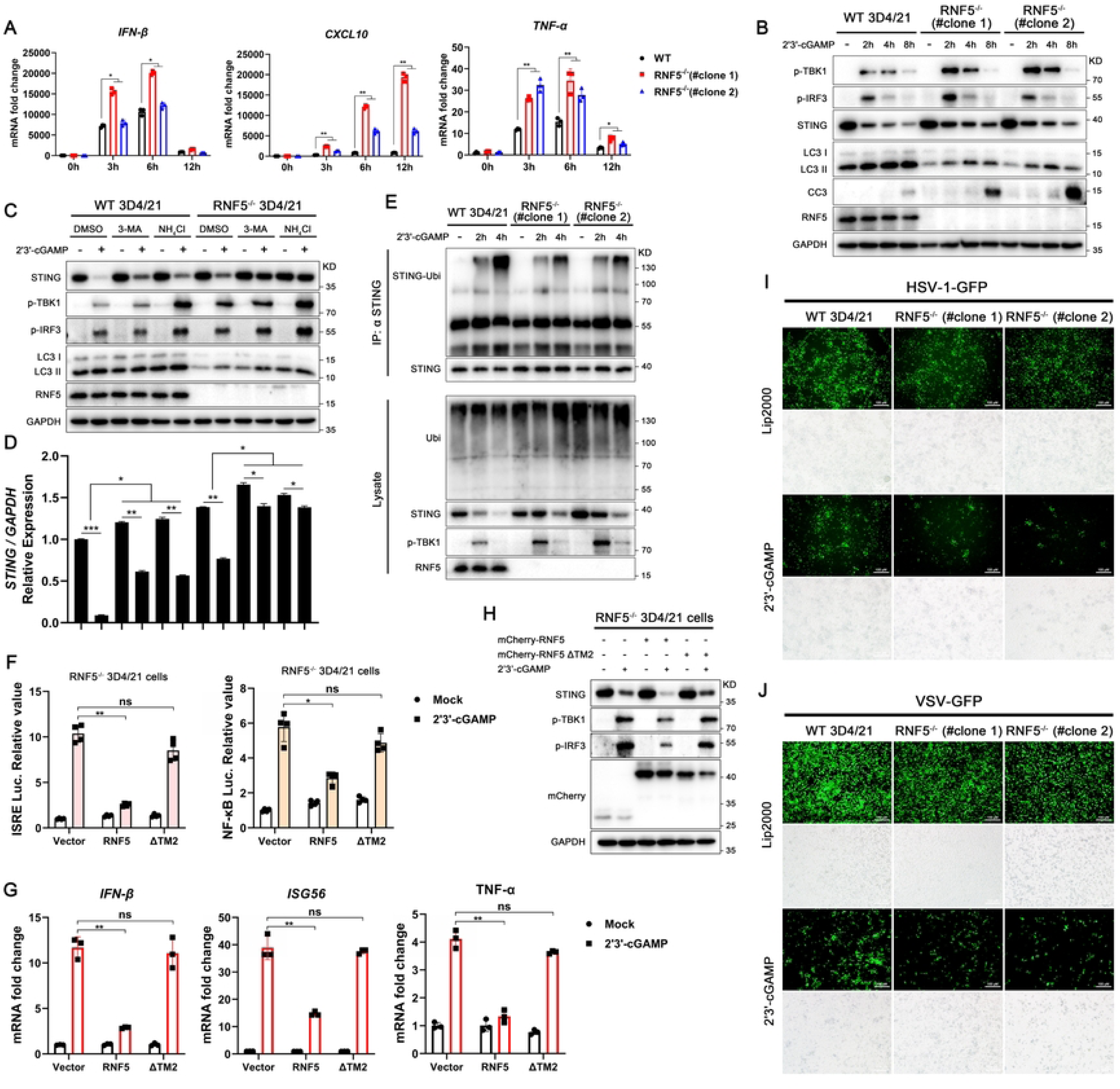
RNF5 negatively regulates pSTING-mediated antiviral response. **(A and B)** RNF5^-/-^ and wild-type 3D4/21 cells were transfected with 2’3’-cGAMP for indicated times, the expression of relevant genes at the transcriptional and protein levels was analyzed by RT-qPCR (A) and Western blotting (B), respectively. **(C and D)** RNF5^-/-^ and wild-type 3D4/21 cells were stimulated by 2’3’-cGAMP (2 μg/mL) for 8 h. Cells were then treated with 3-MA (50 μM), NH4Cl (25 mM), and MG132 (10 μM) for 0.5 h, and analyzed by Western blotting (C). The expression of STING was normalized to GAPDH, and the densitometry values were plotted (D). **(E)** STING was immunoprecipitated from RNF5^-/-^ and wild-type 3D4/21 cells stimulated with 2’3’-cGAMP for varying times, and its ubiquitination level was determined by Western blotting. **(F-H)** RNF5^-/-^ 3D4/21 cells were reconstituted with mCherry-tagged RNF5 or its TM2 deletion mutant. After 24 h, the cells were stimulated with 2’3’-cGAMP for 8 h, then analyzed by luciferase assay (F), RT-qPCR (G), and Western blotting (H). **(I and J)** RNF5^-/-^ and WT 3D4/21 cells were stimulated by 2’3’-cGAMP (2 μg/mL) for 4 h, and then infected with HSV-1 (MOI=0.01, 36 h) (I) or VSV (MOI=0.001, 12 h) (J). Cells were analyzed for viral GFP signal by fluorescence microscopy. \**p* < 0.05, \*\**p* < 0.01 and \*\*\**p* < 0.001.

Complementation of RNF5-knockout 3D4/21 cells with wild-type RNF5 or a TM2-deletion mutant showed that RNF5 complementation markedly suppressed STING activation-induced expression of IFN-stimulated genes and inflammatory genes, as assessed by dual-luciferase reporter assays and quantitative real-time PCR (Figs. 7F and 7G). Mechanistically, RNF5 complementation further promoted STING degradation upon agonist treatment, while the TM2-deletion mutant abolished this effect and restored downstream signaling activation upon STING stimulation (Fig. 7H). Evaluation of the role of porcine RNF5 in regulating cGAS-STING-mediated antiviral signaling revealed that RNF5 deficiency enhances both baseline broad-spectrum antiviral capacity and STING activation-induced antiviral responses upon infection with HSV-1, VSV or PRV (Figs. 7I–7J and S5D-S5I). Viral replication levels and viral titers were significantly lower in RNF5-knockout cells compared to wild-type controls (Figs. 7I–7J and S5D-S5I).

### 2.8. USP20 serves as the primary deubiquitinase that removes K48-linked polyubiquitin chains at K61 of porcine STING

Post-translational modifications of proteins often exhibit dual regulatory effects, both positive and negative. For example, protein kinases mediating phosphorylation and their corresponding phosphatases jointly maintain the homeostasis of protein phosphorylation, thereby balancing intracellular functions. Unlike other post-translational modifications, proteins marked by ubiquitin chains are predominantly directed toward three pathways: (1) recognition and degradation by the 26S proteasome[32], (2) recognition by autophagy receptors and transport to autolysosomes for degradation[33], or (3) removal of ubiquitin chains by deubiquitinases (DUBs) after performing specific functions, allowing cellular recycling[34].

Compared to E3 ligases, STING is regulated by a diverse array of DUBs, most of which are ubiquitin-specific proteases (USPs), including USP20, USP44, USP49, among others[35–40]. In pigs, based on reports from human and mouse studies, we cloned and constructed eight proteases in pigs that directly or indirectly influence STING ubiquitination. Our results showed that multiple proteins, including porcine USP20, USP35, CYLD, EIF3F, and RNF26, significantly removed STING ubiquitination. Among them, USP20 and RNF26 almost completely eliminated RNF5-mediated polyubiquitination of STING (Figs. 8A and 8B). These two proteins negatively regulate STING post-translational modification through distinct mechanisms: RNF26 inhibits STING ubiquitination by reducing RNF5 expression and suppressing its interaction with STING (Fig. 8B), while USP20 directly removes K48-linked polyubiquitin chains from STING regardless of RNF5 presence (Fig. 8B). Even when only the K61 site of STING was retained, USP20 still exhibited complete de-ubiquitination activity (Fig. S6A).

**Figure 8.**
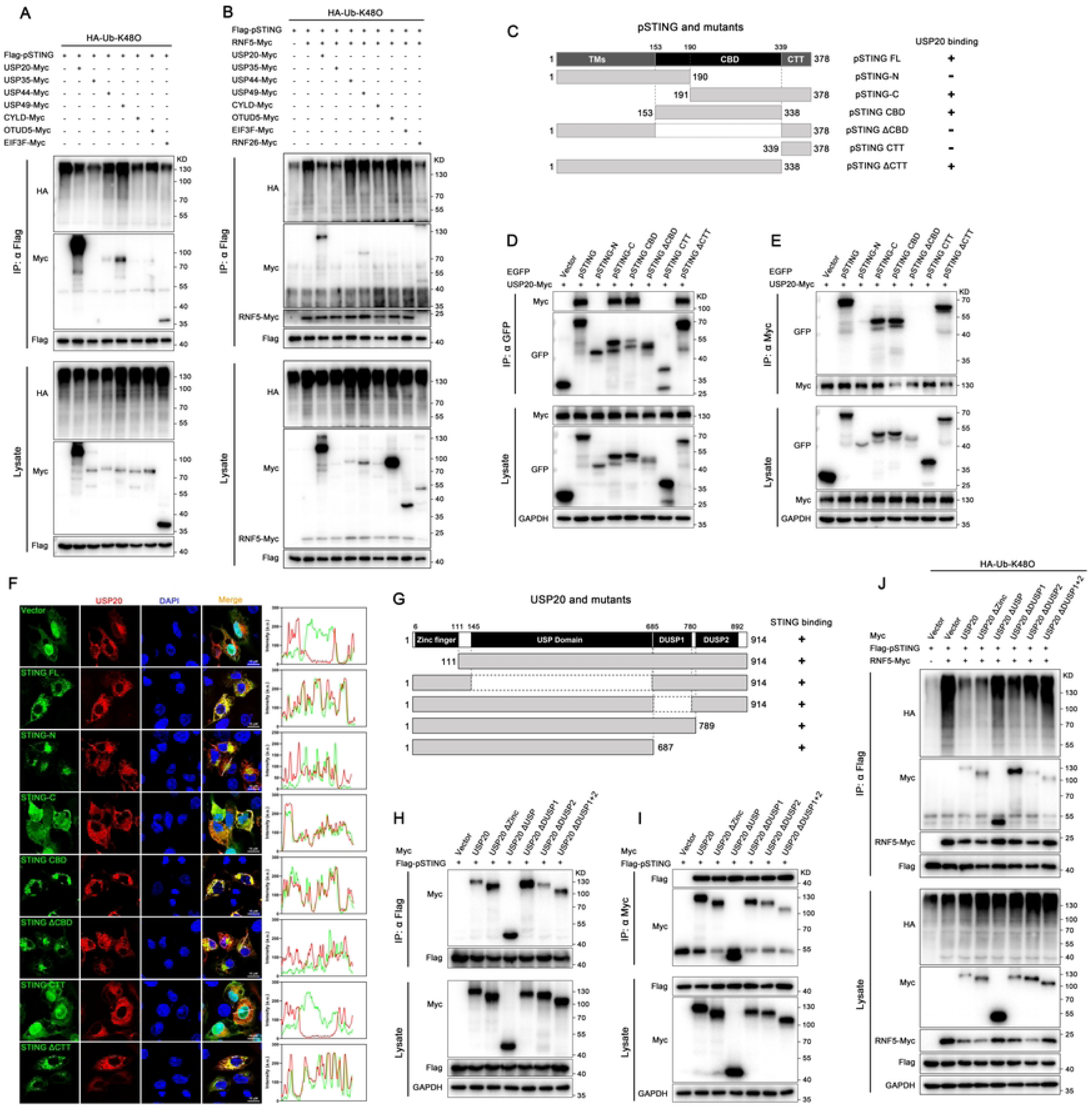
USP20 serves as the primary deubiquitinase that removes K48-linked polyubiquitination at K61 of pSTING. **(A and B)** Under conditions of the absence (A) or presence (B) of pRNF5, co-immunoprecipitation was performed to detect the interaction between Flag-tagged STING and several Myc-tagged deubiquitinases, as well as to determine the level of K48-linked ubiquitination on pSTING. **(C-E)** A schematic diagram of pSTING mutants (C), and reciprocal co-immunoprecipitation analysis of the interaction between pSTING mutants and USP20 in HEK293T cells co-transfected with EGFP-STING mutants and USP20-Myc (D and E). **(F)** The colocalization of pSTING mutants with USP20 was observed by confocal microscopy **(G-I)** A schematic diagram of USP20 mutants (G), and reciprocal co-immunoprecipitation analysis of the interaction between USP20 mutants and pSTING in HEK293T cells co-transfected with USP20-Myc mutants and Flag-pSTING (H and I) **(J)** The effects of each USP20 mutants on the level of RNF5-assembled K48-linked ubiquitin chains on STING was also determined.

We further investigated the key domains mediating the interaction between STING and USP20. Similar to RNF5, deletion of the CBD domain of STING significantly inhibited its interaction with USP20 (Figs. 8C–8E). Subcellular localization analysis showed that loss of the STING CBD domain markedly reduced co-localization with USP20 (Fig. 8F). As a member of the ubiquitin-specific protease family, USP20 contains a canonical USP domain. In addition to the central USP domain, USP20 has an N-terminal zinc finger domain and two C-terminal DUSP domains. To determine which domain of USP20 is critical for interacting with and negatively regulating STING ubiquitination, we generated five domain deletion mutants of USP20 (Fig. 8G). We found that individual deletion of any single domain did not completely abolish the interaction with pSTING, but loss of the DUSP2 domain appears to slightly reduce USP20 binding to pSTING (Figs. 8H and 8I). Furthermore, deletion of the USP domain altered the subcellular localization of USP20 to whole cells, but not altered its co-localization with STING (Fig. S6B). Meanwhile, loss of either the DUSP2 or USP domain greatly impaired USP20-mediated de-ubiquitination of STING (Figs. 8J and S6C). As an E3 ubiquitin ligase and a deubiquitinating enzyme for pSTING, respectively, porcine RNF5 and USP20 regulate pSTING ubiquitination and thereby influence its protein stability. We further analyzed the interactions among these three proteins and found that RNF5 interacts with USP20 even in the absence of STING, and the presence of STING enhances this interaction (Fig. S6D). This suggested that these three proteins form a trimeric complex that regulates the homeostasis of STING protein and downstream signaling.

We further generated USP20-knockout 3D4/21 cells. Although no commercially available antibody targeting porcine USP20 was identified, genetic analysis confirmed the successful disruption of the original open reading frame (ORF) of USP20, yielding multiple homozygous USP20-knockout clones (Fig. S1F and S1G). In contrast to the effects of RNF5 depletion, USP20 deficiency significantly suppressed multiple signaling events following STING agonist stimulation, including the phosphorylation of TBK1 and IRF3, as well as the lipidation of LC3 (Fig. 9A). It also differentially reduced the subsequent transcription of IFN-β, IFN-stimulated genes (ISGs) and TNF-α (Fig. 9B). Mechanistically, USP20 deletion enhanced pSTING ubiquitination and promoted its degradation after activation (Figs. 9C), thereby impairing multiple downstream signals mediated by STING. Furthermore, loss of USP20 attenuated the broad antiviral effects of STING, including against HSV-1, VSV and PRV (Figs. 9D–9H and S6E–S6G).

**Figure 9.**
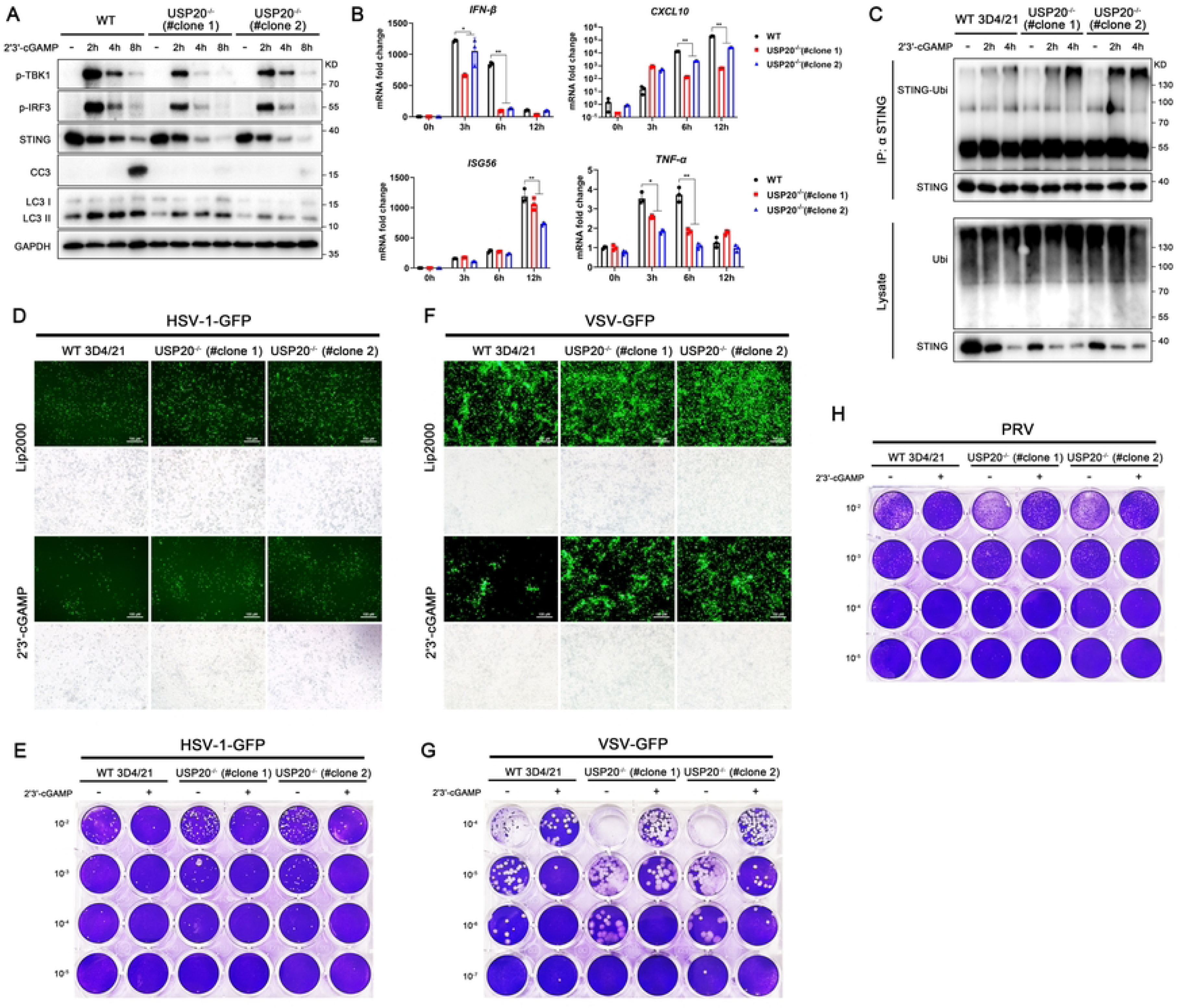
USP20 positively regulates pSTING-mediated antiviral response. **(A and B)** USP20^-/-^ and wild-type 3D4/21 cells were transfected with 2’3’-cGAMP transfection for indicated times, the expressions of relevant proteins and genes were analyzed by Western blot (A) and RT-qPCR (B), respectively. **(C)** STING was immunoprecipitated from USP20^-/-^ and wild-type 3D4/21 cells stimulated with 2’3’-cGAMP for varying times, and its ubiquitination level was determined by Western blotting. **(D-H)** USP20^-/-^ and WT 3D4/21 cells were stimulated by 2’3’-cGAMP (1 μg/mL) for 4 h. and then infected with HSV-1 (MOI=0.01, 36 h) (D and E), VSV (MOI=0.001, 12 h) (F and G) or PRV (MOI=0.01, 24 h) (H). Cells were visualized for viral GFP signal by fluorescence microscopy and viral titers were measured by plaque assay. \**p* < 0.05 and \*\**p* < 0.01.

### 2.9. Ubiquitination at the K61 is not involved in pSTING degradation mediated by selective autophagy

The ubiquitination of cargo proteins is not only essential for proteasome-mediated degradation but is also involved in the process of selective autophagy. Unlike cGAS and TBK1, which are recognized by autophagy receptors via K27-linked ubiquitin chains[41, 42], for STING, autophagy receptors in humans and mice typically recognize through K63-linked ubiquitin chains[13, 18]. However, similar mechanisms have not yet been defined in pigs. Given that various types of polyubiquitination occur on the K61 of pSTING, we sought to investigate whether K61 is also involved in the selective autophagy-mediated degradation of STING. Treatment with various inhibitors revealed that MG132, but not 3-MA or chloroquine, was sufficient to completely reverse the differential expression of pSTING and the K61R mutant (Fig. S7A). We further constructed eukaryotic expression plasmids for multiple autophagy receptors involved in regulating the cGAS-STING pathway, including p62, Tollip, NCP1, CCDC50, and UXT, and investigated their effects on STING protein stability. By co-expressing pSTING with each porcine autophagy receptors in 293T cells, the results showed that although each receptors reduced STING protein expression to varying degrees (Fig. S7B), the K61R mutation of pSTING did not alter the impact of any autophagy receptor on STING stability (Fig. S7C). Co-immunoprecipitation results further indicated that in pigs, the K61R mutation does not affect the interaction between STING and the previously reported autophagy receptors (Fig. S7D). Together, these results demonstrated that the K61 site of pSTING is not involved in selective autophagy-mediated STING degradation.

## 3. Discussion

In this study, we identified the species-specific K61 of porcine STING for K48-linked ubiquitination, regulating STING proteasomal degradation and antiviral activity. Although the regulation of cGAS-STING signaling has been extensively studied in humans and mice, it has not been well defined in pigs. Previous studies have often focused on the functional exploration of conserved sites in certain proteins, with the aim of revealing a common regulatory mechanism across different species[43]; however, species diversity dictates that species-specific regulation is often the direction of evolutionary selection[44]. The activation of NF-κB and autophagy by STING is a relatively ancient and conserved function, yet the regulatory mechanisms differ among species[11, 45]. Notably, the amino acid sequence homology of STING across species is not high; especially, livestock share less than 80% homology with other mammals, and have fewer lysine (K) sites for potential ubiquitination modification compared to primates or rodents (Fig. 4A). The post-translational modifications of the STING, particularly ubiquitination, was reported to vary among species[46, 47].

Unlike E1 and E2 enzymes, which are highly conserved across species[48], the homology of E3 ligases decreases as evolutionary distance increases[49]. The E3 ligases mediating STING ubiquitination exhibit species specificity. For instance, in mice, TRIM30a promotes K48-linked polyubiquitination of STING at K275[50], but humans and pigs lack a fully homologous ortholog of TRIM30a. Furthermore, numerous studies using mouse models have demonstrated the important negative regulatory role of TRIM29 in STING-mediated antiviral immunity[15, 47, 51]. However, it remains unclear whether this effect extends to other mammals. At least in pigs, our results confirm that TRIM29 has minimal impact on the K48-linked polyubiquitination of STING (Fig 5C). RNF5, as a functionally conserved E3 ligase, has been extensively reported to assemble K48-linked polyubiquitin chains on the homologous K150 site of STING in humans, mice, and fish[16, 46, 52]. However, our work in pigs reveals that the primary modification site is K61 rather than K150 (Fig 3C). This indicates distinct site preferences in the degradation mechanisms across different species.

Lysine 61 is highly conserved in mammalian STING; however, the K48-linked ubiquitination was not observed in species such as human, mouse, monkey, and cattle, except in pigs (Figs. 4E–4I). In cattle, which also belongs to livestock, mutating the two amino acids adjacent to K61 (I60 and I63) to the porcine counterparts V60 and L63 heightened K48-linked ubiquitination. Subsequent mutation of K61R on this basis resulted in a phenotype similar to that of porcine STING, where the K61R mutation substantially reduced K48-linked ubiquitination of the bovine STING mutant (Figs. S3D-S3F). This indicated that although lysine is the key amino acid for ubiquitin conjugation, the flanking amino acid background may also be a potential factor influencing ubiquitin chain assembly.

We elucidated two mechanisms of porcine STING degradation, autophagy pathway and particularly ubiquitin-proteasome pathway. RNF5 was found, as an E3 ligase, to bind the CBD domain of pSTING (Figs. 6B–6D) and specifically assemble K48-linked ubiquitin chains at K61 of pSTING, mediating its degradation by the proteasome to prevent excessive STING activation (Fig. 5). This process occurs after STING translocate from the endoplasmic reticulum to the Golgi apparatus and does not affect the recruitment and activation of TBK1 and IRF3 by STING (Figs. 2C, 2D, and 2G). Furthermore, the E3 ligase function of RNF5 depends on its own second transmembrane domain (TM2). The absence of TM2 not only impairs its interaction with STING (Figs. 6F–6H) but also abolishes its regulation of STING stability and subsequent signaling pathways (Figs. 7F–7H and S5C). In contrast to RNF5, USP20, as the primary deubiquitinating enzyme, removes K48-linked ubiquitin chains from the K61 of STING, thereby maintaining the hemostasis of pSTING protein stability and its downstream antiviral signaling (Fig. 9).

Autophagy receptors recognize the ubiquitin chains on cargo proteins through their specific domains, thereby transporting the target proteins to autophagosomes for degradation[53]. We investigated whether STING K61 is involved in the selective autophagy-mediated degradation of STING. However, the results showed that the K61 mutation did not affect the binding of STING to relevant autophagy receptors (Fig. S7D). Previous reports have indicated that K63-linked ubiquitination of STING is a key event recognized by autophagy receptors. In contrast, pSTING primarily undergoes K48-, K27-, K29-, and K33-linked polyubiquitination at K61 (Figs. 3A-D and S3A-C). The K61R mutation does not affect K63-linked ubiquitination of STING (Fig. 3A), likely explaining why this site does not influence autophagy-mediated STING degradation. In our previous study, we identified two conserved LC3-interacting regions (LIR) in pSTING that are involved in regulating ATG5- and ATG16L-dependent autophagy[54]. This represents one of the primary mechanisms of negative feedback regulation in pSTING signaling and does not strictly depend on K63-linked ubiquitination or binding to autophagy receptors[55, 56].

Our data demonstrate that the protein stability of STING impacts on the intensity of multiple downstream signaling pathways (Fig. 2). The mutation of K61 in pSTING enhances various downstream antiviral signals, including IFN, NF-κB, autophagy, and apoptosis, suggesting that the K61 genotype may serve as a molecular marker and target for breeding of antiviral pigs. This idea aligns well with strategies in agricultural breeding that leverage polymorphisms in innate immune genes[57, 58]. Additionally, identification of the specific E3 ligase RNF5 and deubiquitinase USP20 provides potential targets for design of small-molecular antiviral drugs such as RNF5 inhibitors[59], for controlling pig viral diseases.

## 4. Conclusion

we identify here a previously unrecognized and species-specific K61 site in porcine STING that specifically mediates K48-linked polyubiquitination and subsequent degradation of the protein, thereby influencing downstream multiple signaling and antiviral activity (Fig. 10). Although this K61 site is widely present in various mammals and even avian species, it exhibits the specific phenotype only in pigs. Furthermore, we reveal the specific E3 ubiquitin ligase RNF5 and the deubiquitinating enzyme USP20 as regulators of modification at this site, which act antagonistically to maintain the hemostasis of STING activation upon virial infection (Fig. 10). Our work underscores the species-specific fine regulation of innate immunity in pigs, but also provides targets for breeding of antiviral pigs and design of antiviral drugs for pigs.

**Figure 10.**
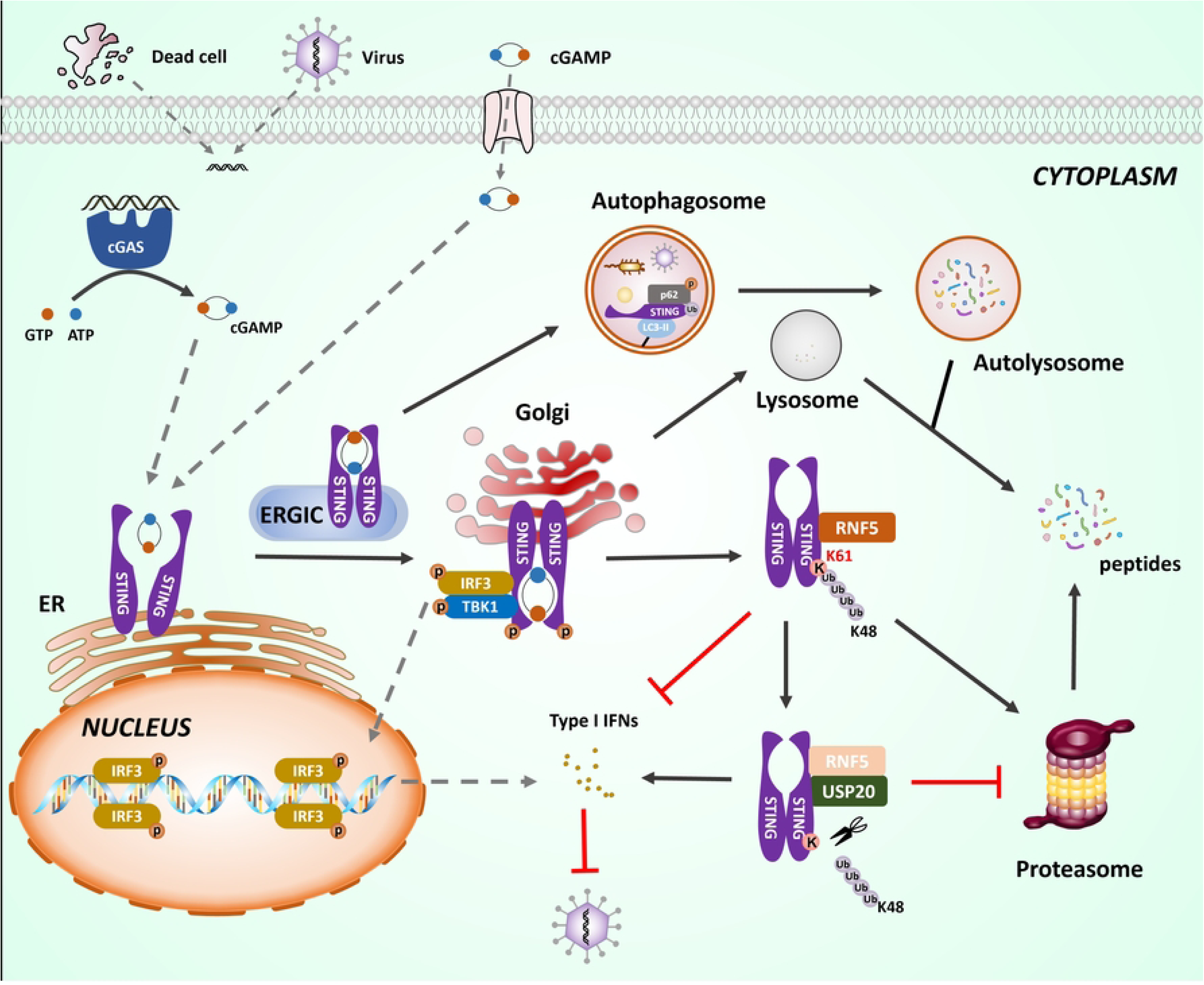
Schematic diagram of the fine regulatory mechanism underlying pSTING degradation and antiviral activity. Upon recognizing 2’3’-cGAMP, either synthesized by cGAS or directly entering the cells, pSTING translocates from the endoplasmic reticulum (ER) to the Golgi apparatus. During this trafficking, it recruits TBK1 and IRF3, thereby activating the IFN response and NF-κB signaling, exerting the antiviral function. Following its activation, STING is subjected to degradation via two distinct pathways: one involves its delivery to lysosomes via autophagosomes or recycling endosomes, while the other entails recruitment of the specific E3 ligase RNF5, which assembles a K48-linked ubiquitin chain at lysine 61 (K61) of pSTING, leading to its recognition and degradation by the proteasome. USP20, regardless of the presence or absence of RNF5, can be recruited by activated pSTING and removes the polyubiquitin chains as the deubiquitinase, thereby maintaining STING stability. The K61, RNF5 and USP20 all play important regulatory roles in pSTING mediated antiviral function.

## 5. Materials and methods

### 5.1. Reagents and antibodies

TRIpure reagent for RNA extraction was obtained from Aidlab (Beijing, China). Double-luciferase reporter assay kits, 2×Taq Master Mix (Dye plus), 2 × Phanta Max Master Mix (P515-01), HiScript® 1st Strand cDNA Synthesis Kit, ChamQ Universal SYBR qPCR Master Mix and 180 kD prestained protein marker were all from Vazyme Biotech Co., Ltd (Nanjing, China). The 2 × MultiF Seamless Assembly Mix was acquired from Abclonal (Wuhan, China). Poly I:C-LMW, 2’3’-cGAMP, and poly dA:dT were purchased from InvivoGen (Hong Kong, China). Golden Star T6 Super PCR mix polymerase was purchased from Tsingke (Nanjing, China) and KOD plus neo polymerase was obtained from Toyobo (Shanghai, China). Annexin V-FITC/propidium iodide (PI) was purchased from Becton Dickinson (Franklin Lakes, NJ, USA). 3-MA (HY-19312), MG132 (HY-13259), NH_4_Cl (HY-Y1269) and Chloroquine (HY-17589AR) were purchased from MedChemExpress (MCE; Shanghai, China). Protein A/G PLUS-Agarose was bought from Santa Cruz Biotechnology (sc-2003, CA, USA). 4’,6-diamidino-2-phenylindole (DAPI) staining solution (C1005), His-tag Protein Purification Kit (P2226) and GST-tag Protein Purification Kit (P2262) were purchased from Beyotime (Shanghai, China). Low melting agarose and CHX (C4859) was purchased from Sigma-Aldrich (St. Louis, USA). All other chemicals and reagents were analytical grade and obtained commercially.

Rabbit mAbs against HA (3724S), LC3I/II (3868), cleaved caspase-3 (Asp175) (9664S), TBK1 (3504S), p-TBK1 (5483S, Ser172), IRF3 (11904S), FLAG (14793), GFP (2956) and β-actin (5057) were acquired from Cell Signaling Technology (Boston, MA, USA). The phosphorylated-IRF3 (Ser396) rabbit mAb (MA5-14947), and goat anti-rabbit IgG (H+L) cross-adsorbed DyLight™ 488 (35553) were obtained from ThermoFisher Scientific (Shanghai, China). The Phospho-STAT1-Y701 Rabbit mAb (AP0054) was acquired from Abclonal (Wuhan, China). The mCherry rabbit pAb (ab183628), goat anti-mouse IgG H&L (Alexa Fluor® 568) (ab175473) and goat anti-rabbit IgG H&L (Alexa Fluor® 568) (ab175471) were acquired from Abcam (Cambridge, UK). The rabbit pAbs of Myc (16286-1-AP), STING (19851-1-AP), ATG5-ATG12L (10181-2-AP), ubiquitin (10201-2-AP) and mouse mAb of GAPDH (60004-1-Ig) were all purchased from ProteinTech (Wuhan, China). The rabbit pAb of RNF5 (abs118428) was obtain from Absin (Shanghai, China). The STAT1 Rabbit mAb (R25799) was purchased from Zenbio (Chengdu, China). Mouse mAbs against HA (HT301-01), GFP (HT801-01), FLAG (HT201-01) and GST (HT601-01) were all acquired from Transgen Biotech (Beijing, China). HRP goat anti-rabbit IgG (H+L) highly cross-adsorbed secondary antibody and goat anti-mouse IgG (H+L) highly cross-adsorbed secondary antibody were all obtained from Sangon Biotech (Shanghai, China). The antibodies against African swine fever virus structural proteins p72 and p30 were prepared and stored by our laboratory.

### 5.2. Cell culture, transfection and viruses

HEK293T cells (ATCC cat # CRL-3216), Vero cells (ATCC cat # CCL-81), Raw264.7 cells (ATCC cat # TIB-71), MDBK cells (ATCC cat # CCL-22) and MA104 cell (ATCC cat # CRL-2378.1) were cultured in DMEM (Hyclone Laboratories, USA) containing 10% fetal bovine serum (FBS, Vazyme Biotech Co., Ltd) and 100 IU/ml of penicillin plus 100 μg/ml streptomycin. Bronchoalveolar lavage derived primary porcine alveolar macrophages (PAMs) and porcine alveolar macrophages (3D4/21, ATCC cat# CRL-2843) were cultured in RPMI (Hyclone Laboratories) containing 10% FBS and penicillin/streptomycin. All the cells were maintained at 37°C with 5% CO_2_ in a humidified incubator. Cell transfection was performed using Lipofectamine 2000 (ThermoFisher Scientific, Shanghai, China) following the manufacturer’s instructions. Herpes simplex virus-1 (HSV-1-GFP), vesicular stomatitis virus (VSV-GFP) and porcine pseudorabies virus (PRV, Bartha K61 strain, GenBank accession JF797217) were stored in our laboratory. The African swine fever virus (ASFV, GenBank accession ON456300) and ASFV-GFP (MGF110-1R gene deleted strain) was preserved in the animal biosafety level 3 (ABSL-3) laboratory of Yangzhou University.

### 5.3. Plasmids and molecular cloning

The FLAG-tagged pCMV plasmid pSTING, the pEGFP -C1 plasmid STING and mutants, the pmCherry-C1 plasmid STING and the HA-tagged pcDNA DEST plasmids pcGAS were previously constructed and used in our laboratory[60, 61]. Total RNA was extracted from Vero cells, Raw264.7 cells, MDBK cells and 3D4/21 cells using TRIpure reagent, and the open reading frames (ORFs) of monkey (mo) STING (XM_008014636), mouse (m) STING (XM_017317994), bovine (b) STING (NM_001046357), porcine (p) RNF5 (NM_001123224), pTRIM29 (XM_021062976), pTRIM13 (XM_021065576), pTRIM7 (XM_021083408), pUSP20 (XM_021070139), pUSP35 (XM_013979184), pUSP44 (XM_021078961), pUSP49 (XM_021098683), pCYLD (XM_021094185), pOTUD5 (XM_021080668), pEIF3F (XM_021062272), sRNF26 (XM_005667432), pNPC1 (NM_214322), pSQSTM1 (XM_005654936), pCCDC50 (XM_003483296), pTollip (NM_001315800), pUXT (XM_003135055), pUBA1 (XM_021080354), pUBE2D1 (NM_001078674) were amplified by RT-PCR using the designed primers as shown in Table S1. The PCR products were cloned into the *EcoR* I and *Sal* I sites of p3×Flag-CMV-7.1 vector, the *EcoR* I and *EcoR* V sites of pCAGGS-Myc/2HA vector, the *Bgl* II and *EcoR* I sites of pmCherry-C1 vector, the *EcoR* I and *Xho* I sites of pET32a-6His vector and the *Xho* I and *EcoR* I sites of pCold I-GST vector, respectively. The Ub-K48 was PCR-amplified from the purchased plasmid pRK5-6×His-Ubiquitin-K48 (Miaolingbio, P37539) and the amplified product was ligated into the pET32a vector to construct prokaryotic expression plasmid. For the construction of the Ub_4_-K48 expression plasmid, the Ub-K48 fragment was first cloned from the template pET32a-6His-Ub-K48, and then the two fragments were linked with an AAA base pair linker. The Ub_2_-K48 was then ligated into the pET32a vector using homologous recombination. Next, Ub_2_-K48 was amplified, and the fragments were joined with an AAG base pair linker to generate the Ub_4_-K48 prokaryotic expression plasmid. All PCR products were inserted into the vector plasmid via homologous recombination using 2 × MultiF Seamless Assembly Mix (Abclonal). The mutation PCR primers for STING of multiple species were designed using the QuickChange Primer Design method (https://www.agilent.com) and are shown in Table S1. Mutation PCR was performed using KOD plus neo polymerase and p3×Flag-CMV-p/mo/m/b/chSTING or pmCherry-C1-pSTING as the template. The PCR products were transformed into competent DMT *E.coli* after *Dpn* I digestion, and the resultant mutants were sequenced. The various ubiquitin eukaryotic expression plasmids were kindly provided by Prof. Peirong Jiao from South China Agricultural University.

### 5.4. CRISPR guide RNA-mediated gene knockout macrophage cell clones

ATG5^-/+^ and ATG5^-/-^ 3D4/21 were previously constructed by us and used in this study[54] (Fig S1A and B). For monkey STING, porcine RNF5 and porcine USP20 gene knockout (KO), CRISPR gRNAs were designed and the KO cell clones were obtained, as we described previously[54, 62, 63]. The DNA sequences encoding all gRNAs are shown in Table. S2. Single clones of MA104 cells or 3D4/21 cells were used to detect protein expression of STING or RNF5 by Western blotting. All MA104 and 3D4/21 cell clones were subjected for detection of genomic DNA editing by sequencing the PCR products, as described[54, 62, 63], using the PCR primers shown in Table S2. Homologous KO cell clones were obtained for RNF5 and USP20, respectively (Fig. S1C-G).

### 5.5. Promoter-driven luciferase reporter gene assay

The dual luciferase reporter assay was performed as we described previously [64]. The results were expressed as fold induction of ISRE or ELAM (NF-κB)-Fluc expression, relative to that of the control vector, following Fluc normalization by the corresponding Rluc values.

### 5.6. RNA extraction and reverse transcription-quantitative PCR

The RNA extraction, reverse transcription and quantitative PCR were performed as we described previously[54]. The sequences of the qPCR primers used in this study are listed in Table S3.

### 5.7. Western blotting and Co-immunoprecipitation

Whole cell protein extraction, protein sample processing and Western blotting were performed as we described previously [54, 64, 65]. HEK293T cells in 6-well plates (8×10^5^ cells/ well) were transfected with different combinations of plasmids for 24 h. Then, the cells were harvested and lysed in 300 μL radioimmunoprecipitatation assay (RIPA, 50 mM Tris-base, pH 7.5, 150 mM NaCl, 0.5% Triton X-100, 0.5% sodium deoxycholate, and 1 mM PMSF) buffer containing protease inhibitors on ice for 30 min.

The cell lysates were centrifuged at 12,000 ⅹ g to pellet the cellular debris, and the cleared supernatants were used for co-immunoprecipitation as we described previously [64].

### 5.8. Cellular ubiquitination assay

HEK293T cells, 3D4/21 cells, or PAMs were transfected with specified constructs or infected with different viruses. Prior to collection, the cells were treated with 10 μM MG132 (MCE) for 6 h. After the indicated durations, cells were harvested. Following PBS washing, the cells were lysed on ice for 10 min using RIPA buffer, with 10 mM N-ethylmaleimide. The lysates were sonicated and centrifuged at 12,000 × g for 10 min at 4°C. The supernatant was incubated and immunoprecipitated with specific antibodies overnight at 4°C. Protein A/G agarose beads were then added and incubated at 4°C for 4 h. After extensive washing, the bound proteins were eluted with 2× SDS sample buffer and analyzed by immunoblotting.

### 5.9. Protein purification

Wild-type porcine STING and the K61R mutant were cloned into a pCold I vector with an N-terminal GST tag, while porcine UBA1, UBE2D1, RNF5, along with single Ub-K48 ubiquitin or a tandem chain of four Ub-K48 ubiquitin, were cloned into a pET32a vector with an N-terminal 6×His tag. For protein expression, a single colony of transformed *E. coli* BL21 cells was cultured overnight in LB medium supplemented with 100 µg/mL ampicillin. The culture was then diluted 1:100 into fresh LB medium and grown at 37°C for 2–4 h until the optical density at 600 nm (OD₆₀₀) reached 0.6. After adding IPTG to a final concentration of 0.5 mM, the cells were further cultured overnight at 16°C.

For GST-tagged protein purification, the washed cell pellet was resuspended in lysis buffer (PBS, 1 mg/mL lysozyme). After sonication, the supernatant of the cell lysate was loaded onto a GST-tag affinity chromatography column containing BeyoGold™ GST-tag Purification Resin (Beyotime, P2250). After the beads were washed three times with lysis buffer, the purified protein was then eluted with elution buffer (50 mM Tris-HCl, 150 mM NaCl, pH 8.0) containing 10 mM reduced glutathione (GSH). Alternatively, for interaction studies, the beads were incubated with a specified protein in binding buffer (50 mM Tris-HCl, 200 mM NaCl, 1% NP-40, 1 mM DTT, 1 mM EDTA, 10 mM MgCl₂, 1 mM PMSF, pH 8.0) at 4°C with gentle agitation before elution. For His-tagged protein purification, the washed cell pellet was subjected to sonication in lysis buffer (PBS, 1 mg/mL lysozyme, pH 8.0). The fusion protein was then immunoprecipitated using nickel-NTA beads. The beads were washed three times with wash buffer (PBS, 2 mM imidazole, pH 8.0), and the target protein was eluted with elution buffer (PBS, 50 mM imidazole, pH 8.0). The eluted protein was subsequently dialyzed and concentrated. The purity and integrity of all purified proteins were verified by SDS-PAGE, and the proteins were stored at -80°C.

### 5.10. GST-pulldown assay

For the GST pull-down assay, BeyoGold™ GST-tag Purification Resin was coated with 10 μg of either GST-fused STING protein or GST control in binding buffer (50 mM Tris-HCl, 200 mM NaCl, 1% NP-40, 1 mM DTT, 1 mM EDTA, 10 mM MgCl₂, pH 8.0) supplemented with 1 mM PMSF for 2 h at 4°C. The coated beads were then incubated with specified amounts of RNF5 soluble recombinant protein overnight at 4°C following three washes. After incubation, the beads were washed three times, and the bound proteins were analyzed by Western blotting.

### 5.11. *In vitro* ubiquitination assay

Recombinant STING protein with N-terminal GST was incubated with active recombinant E1 (UBA1), E2 (UBE2D1), E3 (RNF5), Ub-K48 or Ub_4_-K48 and ATP (200 mM in 20 mM HEPES pH 7.4, 100 mM magnesium chloride) in ubiquitination assay buffer (20 mM HEPES pH 7.3, 2 mM magnesium chloride, 1 mM EDTA, 1 mM TCEP, 0.05% Tween-20 and 1 mM PMSF). Mixtures omitting ATP were included as negative controls. Alternatively, recombinant GST-STING and GST-STING K61R proteins were incubated the lysates of ubiquitin transfected cells. Reactions were incubated at 37 °C for 2 h and quenched by incubating at 95 °C for 10 min, followed by Western blotting analysis.

### 5.12. Fluorescence microscopy

3D4/21 cells grown on a 15 mm glass bottom cell culture dish (NEST, Wuxi, China) were transfected with indicated plasmids using Lipofectamine 2000. Twenty-four hours later, the cells were fixed with 4% paraformaldehyde for 30 min and permeabilized with PBS (0.5% Triton X-100) at room temperature for 20 min. After washing with PBS, the cells were sequentially incubated with primary Flag rabbit/mouse mAb (1:200) or Myc rabbit pAb (1:200), followed by incubation with secondary goat anti-rabbit IgG DyLight™ 488 (1:800) or goat anti-rabbit/mouse IgG H&L (Alexa Fluor® 568). The stained cells were counter-stained for cell nucleus with 0.5 mg/mL DAPI (Beyotime, Shanghai China) at RT for 15 min. Finally, the cells were visualized under a laser-scanning confocal microscope (LSCM, Leica SP8, Solms, Germany) at excitation wavelengths of 405 nm and 488 nm.

### 5.13. Flow cytometry for detection of apoptosis

The level of cell apoptosis was examined using the Annexin V-fluorescein isothiocyanate (FITC) apoptosis detection kit (Franklin Lakes). Briefly, after treatment, the cells were harvested and washed with the binding buffer, and then resuspended in the binding buffer. The staining solutions of Annexin V-FITC and PI were added one by one. The cells were incubated at RT for 30 min in the dark, and the stained cells immediately detected using flow cytometry. The ratios of Annexin positive cells including PI negative and positive were calculated as the levels of apoptotic cells.

### 5.14. Viral infections and titrations of viral replications

293T, 3D4/21 or MA104 cells were first transfected with the indicated plasmids for 24 h or stimulated with 2’3’-cGAMP for 12 h. Subsequently, the cells were infected with different viruses for various time periods: HSV-1 for 36 h, ASFV for 48 h, PRV for 24 h and VSV for 12 h. Viral titers were then measured by a standard plaque assay in Vero cells for HSV-1, PRV and VSV, which was performed as we described previously[64, 65]. Plaque formation would take 1-4 d depending on the viruses being analyzed (HSV-1 for 3 d, VSV and PRV for 2 d). After staining, Vero cells were washed with tap water until clear plaques appeared. The plaques were counted, and photos were taken. Alternatively, Viral replications were visualized for GFP expression by fluorescence microscope (HSV-1-GFP, ASFV-GFP, VSV-GFP), measured by flow cytometry for GFP positive infected cells (ASFV-GFP), by quantitative qPCR for viral genes of HSV-1 (gD), ASFV (B646L and I267L) and PRV (gD), by Western blotting detection of ASFV structural proteins (p72 and p30).

### 5.15. Quantification and statistical analysis

Data were analyzed using GraphPad Prism 9.5.0 software (GraphPad Software, Inc) and are presented as mean ± SEM. Statistical analysis was performed using unpaired *t* tests or *ANOVA* plus *Least Significant Difference (LSD)* for multiple comparisons. The *p* values were calculated and statistics are represented as follow: NS, not significant (*p* > 0.05); * *p* < 0.05; ** *p* < 0.01 and *** *p* < 0.001. Statistical significance was set at *p* ≤ 0.05.

## Author contribution statement

JZ.Z and Wanglong.Z conceived and designed the experiments; N.X, Y.C, C.C, Z.S, A.L, Wangli.Z, J.J, H.H, J.H, and JJ.Z performed the experiments; N.C, S.J and JZ.Z analyzed the data; N.X and JZ.Z wrote the paper. All authors have contributed to the manuscript and approved the submitted version.

## Data Availability Statement

The raw data are all available upon request.

## Acknowledgements

This work was partly supported by the National Natural Science Foundation of China (32473040; 32202818; 32172867), 111 Project under Grant D18007 and A Project Funded by the Priority Academic Program Development of Jiangsu Higher Education Institutions (PAPD). N.X was supported by the Postgraduate Research & Practice Innovation Program of Jiangsu Province (KYCX23_3596).

